# WISER: spectral whitening of fixed effects for improved breeding value estimation in structured, unbalanced, and genotype–environment-confounded populations

**DOI:** 10.1101/2025.07.17.665325

**Authors:** Laval Jacquin, Walter Guerra, Mariusz Lewandowski, Andrea Patocchi, Marijn Rymenants, Charles-Eric Durel, François Laurens, Maria José Aranzana, Lidia Lozano, Tristan Mary-Huard, Hélène Muranty

## Abstract

This work introduces WISER (whitening and successive least squares estimation refinement), a whitening-based method designed to improve breeding value (BV) estimation by accounting for population structure and genetic covariance within experimental designs. WISER provides a generalized framework applicable across diverse experimental designs, species, and omics datasets, including single nucleotide polymorphisms (SNPs), metabolomics, and near-infrared spectroscopy (NIRS) used as phenomic predictors. Central to WISER is whitening, a statistical transformation that decorrelates variables and standardizes their variances. From a spectral perspective, WISER reweights the fixed-effect design according to the eigenspectrum of the genetic covariance matrix, attenuating dominant directions of genetic variation rather than explicitly correcting for a predefined subset of eigenvectors. This transformation aims to reduce confounding between genetic and environmental effects during BV estimation. In simulations with known genetic values, WISER outperformed BLUE in breeding value (BV) estimation accuracy as genotype-by-environment confounding increased, while showing similar performance under low confounding. Across four species and 33 traits, genomic predictive ability using WISER-estimated BVs was generally higher than with LS-means or BLUP. Because predictive ability depends on the estimation method, we validated this metric via simulation, confirming that it never exceeded the accuracy achieved using true genetic values. These results support WISER as a flexible framework for BV estimation and downstream omics-based prediction in structured and unbalanced experimental populations. A user-friendly R package, *wiser*, has been developed to facilitate its application.

## 1. Introduction

In modern quantitative genetics, accurate breeding value (BV) estimation relies on statistical models that capture phenotypic variation across diverse environments. A major challenge arises when population structure—whether from genetically distinct subgroups or continuous relatedness—is not properly accounted for. This can confound genetic effects, leading to spurious associations and biased estimates [1–5]. Correcting for population structure is therefore essential in genomic selection and GWAS [6–12].

When phenotypes are measured once per individual, population structure can be modeled using the classical kinship matrix [13]. Here, we address the case of repeated measures in experimental designs. Various methods have been developed to correct for unequal relatedness, including PCA [10] and linear mixed models [8]. Eigen-information from kinship or genomic covariance matrices has also been used as fixed-effect covariates in GBLUP and its derivatives [6, 7, 11, 14, 15].

Population structure is particularly relevant in plant breeding, where relatedness may reflect geographic origin [16] or breeding practices [17]. Despite its impact, it is often underestimated [18–20]. In two-step approaches, BVs are typically estimated using LS means [19, 21–29], but this ignores genetic covariance and may introduce confounding in unbalanced designs. BLUP [30–33] is another common approach, sometimes with PCA covariates. However, PCA may not reliably capture population structure [34–38], and has been shown to fail in controlling false positives [35, 37] or to reduce true positive detection [35, 39].

To address this need, we introduce WISER (whitening and successive least squares estimation refinement), a novel approach specifically developed to efficiently model the genetic covariance structure among individuals within experimental designs. This approach accounts for genetic covariance within the experimental design during the initial estimation of individual BVs, before downstream genomic prediction or GWAS. It is based on the premise that accounting for genetic covariance directly during BV estimation, rather than only during downstream genomic prediction, can improve the quality of the resulting BVs, as described in [40].

At the core of WISER is whitening, a transformation that removes correlations and standardizes variances [41–44]. This provides a flexible framework applicable across species, experimental designs (e.g., alpha lattice, multi-environment trials), and omics datasets such as SNPs, metabolomics, and NIRS. Its versatility makes WISER especially valuable in breeding, where diverse experimental setups and integrated omics improve marker-trait associations.

WISER extends eigen-information approaches [6, 7, 11, 45] by whitening fixed-effect variables, thereby reweighting the principal directions of genetic variation according to the spectral structure of the genetic covariance matrix. This transformation may attenuate the influence of dominant directions of genetic variation, including those associated with population structure, on fixed-effect estimation, thereby potentially reducing confounding between genetic and environmental effects and improving BV estimation. This is particularly relevant in plant breeding, where accurate omics-based predictions support crop improvement [18–20]. The method is implemented as a user-friendly R package with automated parameter optimization, applicable to both plant and animal datasets, and integrates seamlessly into genomic prediction and GWAS pipelines, enabling multi-omics analyses [46].

The remainder of this article is organized as follows. Sections 2.1–2.2 present the linear mixed model and its connection to WISER. Supplementary File 1 demonstrates how WISER extends several population-structure correction methods based on eigen-information, including the covariance analysis via eigenvectors (EVG) approach [6, 15] and principal component (PC) models, by relaxing their assumptions and introducing a whitening-based framework. Section 2.3 describes four datasets (rice, maize, apple, Scots pine), simulations, predictive models, and heritability estimates used to compare WISER with LS-means and BLUP. Section 3 reports the results, showing improved estimation accuracies and improved genomic predictive abilities for high levels of confounding when using WISER-derived BVs compared with the alternative BV estimation approaches evaluated. The implications for genomic selection and potential extensions to association mapping are discussed in Section 4.

The R package wiser can be easily installed from GitHub and is permanently archived on Zenodo. In accordance with the FAIR principles, all repositories containing the datasets and R scripts used in this study are publicly available on GitHub, with links provided in the Availability of Data and Materials section at the end of this article.

## 2. Materials and methods

### 2.1 The linear mixed model and the impact of whitening fixed-effect covariates

In the following, we will use the term *genotype* exclusively to mean cultivar, accession or animal. Consider the following linear mixed model:

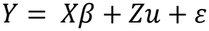

where:

- *Y*(*n* x 1) is the vector of raw phenotypic measurements, with values possibly repeated for each genotype across one or more environments.
- *X*(*n* x *l*) is the fixed-effect design matrix linking the overall mean and *l* − 1 environmental effects to raw phenotypic measurements.
- *β* = (*µ, β*_1_, . ., *β*_*j*_, . ., *β*_*l*−1_ )′ is the vector of fixed effects, comprising the overall mean *µ*and the *l* − 1 fixed environmental effects.
- *u* (*q* x 1) is the vector of random genetic values for *q* genotypes, where 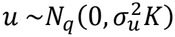 and *K* denotes the genetic covariance matrix between genotypes estimated from omic data.
- *Z*(*n* x *q*) is the design matrix linking genetic values to raw phenotypic measurements in the experimental design.
- *ε* (*n* x 1) is the vector of residuals, where 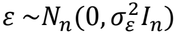.

In this model, the estimation of *u* can be biased if the structure of the experimental design—hereafter referred to as the experimental design population structure or simply population structure— introduces significant confounding between fixed and genetic effects. One common source of such structure is imbalance in the experimental design.

Whitening is a statistical transformation that eliminates correlations between variables and standardizes their variances. In the linear mixed model, a whitening matrix *W* can be derived from a regularized form 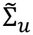 of the genetic covariance structure among individuals, captured by 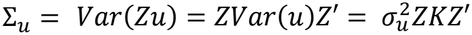. Applying the whitening matrix *W* to the fixed-effect design matrix *X* induces a spectral reweighting of the components of *X* aligned with the eigendirections of the genetic covariance matrix. As detailed in Supplementary File 1, this transformation may attenuate the influence of dominant directions of population structure on fixed effects, potentially reducing confounding between fixed and random effects. Conceptually, WISER plays a role analogous to ridge regression relative to PC-based approaches for population structure correction. Whereas EVG and PC-based methods account for population structure through a finite set of retained eigendirections, WISER preserves the corresponding eigenspace and continuously reweights its components according to the spectral structure of the genetic covariance matrix. In this sense, WISER may be viewed as a softer spectral correction strategy, analogous to the relationship between ridge regression and PC regression.

A whitening matrix *W* satisfies the following condition, known as the whitening property [43, 46, 47]:

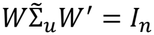

where *I*_*n*_ is the identity matrix of size *n*. This property ensures that the transformed variables are uncorrelated and have unit variance. Importantly, for any full rank covariance matrix, there exists a (generally non-unique) matrix *W* that fulfills this condition.

### 2.2 The WISER statistical framework

Using the same definitions as for the components of the linear mixed model, the WISER statistical framework solves the following model to estimate a vector *v* = ( *v*_1_, …, *v*_*q*_)′ of BVs, which are treated as fixed effects:

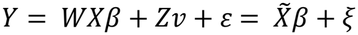

where 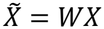 and *ξ* = *Zv* + *ε*. Note that *v* is modeled as a distinct vector of fixed effects relative to *β*, and is not treated as a vector of random effects with a genetic covariance matrix *K* estimated from omic data. This ensures that the estimation of *v* is not derived as a linear or non-linear function of omic effects (e.g., marker effects), as it typically is in genomic BLUP in the linear case. As a result, the vector 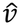 of WISER-estimated BVs can be used in downstream omic-based selection and association analyses. These estimated BVs can be seen as adjusted for confounding between fixed and random effects, providing an adjusted input for genomic prediction and association studies.

The whitening matrix *W* is derived from the regularized genetic covariance matrix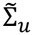. It plays a pivotal role as it transforms the fixed environmental variables to minimize the confounding between environmental and genetic effects. There are several methods for computing a whitening matrix *W*, including zero-phase component correlation analysis (ZCA-cor), principal component correlation analysis (PCA-cor) and Cholesky matrix decomposition (Cholesky) among others [43]. ZCA-cor and PCA-cor whitening matrices are calculated using the spectral decomposition of 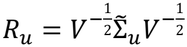, the genetic correlation matrix associated with 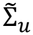, where 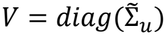 is a diagonal matrix with the variances of 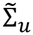 on its diagonal. Since *R*_*u*_ is symmetric and positive semi-definite, it has the following spectral decomposition: 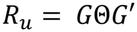, where *G* is the orthogonal eigenvector matrix for *R*_*u*_, while Θ is the diagonal matrix of positive eigenvalues for *R*_*u*_. The inverse square root matrix of *R*_*u*_ is given by 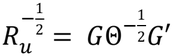, which satisfies 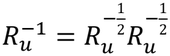 since *G* is an orthogonal matrix. According to [43], the whitening matrices associated to ZCA-cor, PCA-cor and Cholesky are given by:

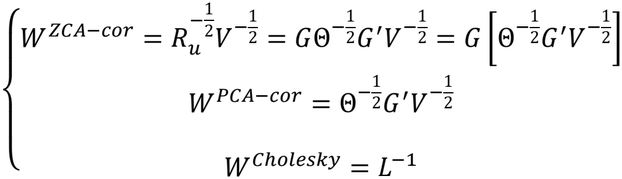

where *L* is a full-rank lower triangular matrix derived from the following Cholesky decomposition 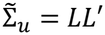 . The PCA-cor whitening procedure can be seen as standardizing variables using 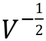, followed by a rotation using the transposed correlation eigenmatrix *G*′, and then scaling using the inverted correlation singular values matrix 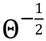. ZCA-cor whitening extends this by applying an additional rotation *G* to revert to the original basis of the standardized variables. While [43] recommends both ZCA-cor and PCA-cor, each whitening method is optimal under distinct criteria. WISER implements these two approaches alongside the Cholesky procedure. Notably, ZCA-cor uniquely ensures that the whitened variables maintain the highest correlation with the original variables, [43] provides a detailed discussion of these criteria and the optimality of each method.

Constructing the whitening matrix matrix *W* depends on the regularized form 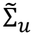 of 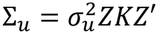, which requires estimating 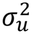. In WISER, 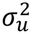 and 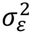 are estimated using a parallelized approximate Bayesian computation (ABC) algorithm for fast and stable variance component estimation in large, complex datasets. Since Σ_*u*_ is rank-deficient in many practical applications, it is often singular and therefore unsuitable for matrix inversion. To overcome this limitation, WISER applies shrinkage and regularization procedures to construct a regularized covariance matrix 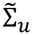 that is full-rank and positive-definite, thereby enabling the computation of whitening matrices [48–53]. Further details are provided in Supplementary File 2.

To automate whitening and regularization choices, WISER includes an optimization function that performs a grid search across whitening methods, parameters, and prediction models (GBLUP, LASSO, SVR, RKHS, random forest). Performance is evaluated via k-fold cross-validation, with the best method chosen by majority vote on mean squared error. By default, optimization is run on a dataset subset (up to 5,000 phenotypic records) for efficiency.

Once the whitening matrix is computed, estimates for *β* and *v* are obtained using the following successive OLS procedure:

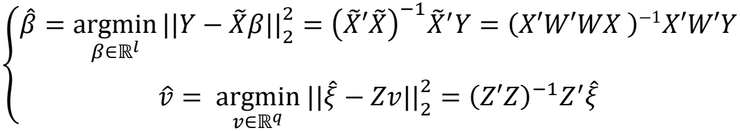

where 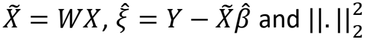 represents the squared *l*_2_ norm. Note that 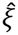 corresponds to the part of *Y* that is not explained by the transformed fixed-effect variables and is orthogonal to these variables. As with any linear model, this orthogonality arises because 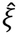 is estimated by projecting *Y* onto the space orthogonal to that spanned by the whitened fixed-effect variables, ensuring orthogonality by design:

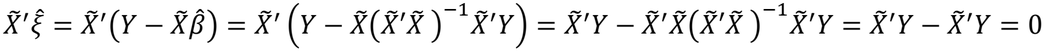

Thus, the vector 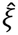 can be used to estimate the BV vector *v*. Importantly, the estimate 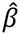 is obtained from the whitened design matrix 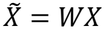, where *W* is derived from the regularized covariance matrix 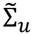. Consequently, the estimation of *β* is informed by the estimated genetic covariance structure and therefore depends, through *W*, on the variance component 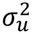. Furthermore, when no repeated phenotypic measurements per genotype are available, the genetic covariance matrix Σ_*u*_ associated with individuals in the experimental design reduces to the genetic covariance matrix *K* associated with genotypes, so that Σ_*u*_ = *K*. To mitigate potential numerical singularities of 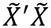 or *Z*^′^*Z* in practical applications, WISER also incorporates the Moore–Penrose generalized pseudo-inverse for the estimation of *β* and *v*.

### 2.3 Analyzed datasets, compared BV estimation methods, predictive models, and computed statistics

#### 2.3.1. Analyzed datasets

Four datasets from experimental designs involving rice [18], maize [20], apple [54], and Scots pine [55] were used to compare genomic predictive ability (PA) and heritability estimation across 33 traits. In these datasets, the omic data were SNP data. The rice dataset contained 4,308 raw phenotypic measurements and 9,928 SNPs associated with 334 genotypes. The 334 rice genotypes originated from a synthetic population developed from 35 tropical japonica (*Oryza sativa* L.) families, following three recombination cycles and advancement to *S*_0:2_ and *S*_0:3_ generations [18]. The maize dataset included 18,970 raw phenotypic measurements and 41,722 SNPs associated with 246 genotypes, corresponding to a diversity panel of dent maize hybrids [20]. The apple reference population (REFPOP) dataset, after preprocessing to remove phenotypic outliers, consisted of 42,522 raw phenotypic measurements and 303,240 SNPs associated with 534 genotypes. These genotypes included both a set of accessions and a set of progenies. The accessions derived from 10 geographical origins, while the progenies belong to 27 distinct families. Among the SNPs, only 50,000 were used, selected through uniform sampling to reduce computation time and resource usage, as [19] demonstrated that a PA plateau is typically reached with as few as 10,000 SNPs for this dataset. Additionally, the apple dataset included two traits—scab and powdery mildew—not covered in [54]. Finally, the Scots pine dataset included 1,248 raw phenotypic measurements and 20,795 SNPs for 208 genotypes from 40 families across 5 native *P. sylvestris* populations, evaluated at two sites [55].

The traits predicted and analyzed for each species were as follows:

- Rice: days to flowering (FL), plant height (PH), grain yield (YLD), and grain zinc concentration (ZN).
- Maize: anthesis, anthesis-silking interval, ear height, grain number, grain weight, grain yield, plant height, silking, and tassel height.
- Apple: color over, flowering start, flowering end, full flowering, flowering intensity, fruit number, single fruit weight, fruit weight, harvest date, powdery mildew, russet frequency, scab, trunk diameter, and trunk increment.
- Scots pine: height (H), annual height increment (I), duration of budburst (D)—the time taken to progress from stage 4 (scales open along the length of the shoot, no needles) to stage 6 (green-tipped needles visible), and time taken to reach individual budburst stages: stage 4 (T4), stage 5 (T5), and stage 6 (T6).

For each dataset, the environmental variable was defined as a combination of available factors such as site (or country), year, management type, block, and genotype generation. In the rice dataset, it was defined by site, year, block, and genotype generation, resulting in 48 unique environments. For maize, it was defined by site, year, management type (rainfed or irrigated), and block, yielding 1,082 levels. In the apple dataset, the environmental variable was defined as the combination of country, year, and management type (standard irrigation and pesticide use, standard irrigation with reduced pesticide use, or reduced irrigation with standard pesticide use), with 75 distinct levels. For pine, the environmental variable was based on site, year, and block, also resulting in 48 levels.

For each dataset, population structure was visually assessed using three complementary approaches applied to the SNP data: a heatmap of the genomic covariance matrix, principal component analysis (PCA), and uniform manifold approximation and projection (UMAP) [56]. These analyses were implemented using the R packages `pheatmap` [57], `mixOmics` [58] and `umap` [59].

Raw phenotypes for a generic trait were also simulated for each dataset using fixed effects, random genetic effects, and random genotype-by-environment interaction effects. The trait heritability was set to a moderate value of 0.5, and the total phenotypic variance was fixed at 1. The simulation procedure was as follows:

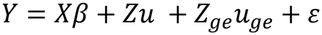

where:

- *Y*(*n* x 1) is the vector of simulated raw phenotypic measurements, with values possibly repeated for each genotype across one or more environments.
- *X*(*n* x *l*) is the design matrix linking fixed effects, the overall mean and the *l* – 1 environmental effects, to raw phenotypic measurements.
- *β* = (*µ, β*, . ., *β*, . ., *β*_′_ is the vector of fixed effects where *µ ∼ U*(0,1) and *β ∼ N*(0,3).
- *u* (*q* x 1) is the vector of random genetic values for *q* genotypes, where 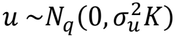 and *K* denotes the genetic covariance matrix between genotypes estimated from omic data.
- *Z*(*n* x *q*) is the design matrix linking genetic values to raw simulated phenotypic values in the experimental design.
- *u*_*ge*_ (*q*(*l* − 1) x 1) is the vector of random genotype by environment interaction effects, where 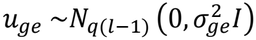 and 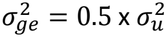.
- *Z*_*ge*_ (*n* x *q*(*l* − 1)) is the design matrix linking random interaction effects to raw simulated phenotypic values in the experimental design.
- *ε* (*n* x 1) is the vector of residuals, where 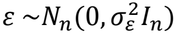.

This simulation framework was used to assess the accuracy of estimating and predicting the simulated genetic values *u*, with and without whitening of the fixed-effect variables, thereby quantifying the impact of the whitening transformation. The effect of model misspecification on the performance of the whitening approach was further evaluated using two scenarios. In the first scenario, the intercept and all environmental effects were included as fixed effects when estimating *u*, corresponding to a correctly specified model. In the second scenario, only the intercept was included as a fixed effect, resulting in a naïve estimate of *u*. For each scenario, 100 phenotype datasets were generated and analyzed.

A third scenario, derived from the first scenario, was designed to investigate the effect of genotype– environment confounding. For each simulation, the experimental design was resampled to achieve a predefined target level of association between genotypes and environments, quantified using Cramér’s *V*. The genomic data was subsequently restricted to the genotypes retained in the resampled experimental design. This procedure generated 100 phenotype datasets, spanning different levels of genotype–environment association, which were subsequently analyzed to evaluate the effect of confounding on estimation performance.

For all three simulation scenarios, ZCA-cor whitening was used, and the diagonal regularization parameter was defined using an empirically calibrated scaling rule (see Supplementary File 2) to maintain computational tractability while preserving a comparable level of regularization across sample sizes.

#### 2.3.2. Compared BV estimation methods

Four BV estimation approaches were compared: LS-means, two BLUP approaches, and WISER.

To account for spatial variation before BV estimation with LS-means and BLUP, spatial heterogeneity correction was applied per trait–environment combination, yielding spatially adjusted phenotypes. This was possible for maize, apple, and pine (using row/column, row/position, and GPS coordinates, respectively), but not for rice, where spatial data were unavailable. Corrections were performed using the spline-based spatial analysis of field trials method [60], as in [19]. In WISER, spatial modeling is detailed below. Residual independence was verified with spatial plots and variograms, which showed no clustering and generally flat patterns, indicating minimal remaining spatial structure.

For the LS-means approach, the following multiple linear regression model was initially fitted:

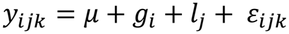

where *y*_*ijk*_ was the *k*^*th*^ spatially adjusted raw phenotype associated to genotype *i* in environment *j, µ* was the fixed overall mean, *g*_*i*_ was the fixed effect of genotype *i* (1 ≤ *i* ≤ *q*), *l*_*j*_ was the fixed effect of the *j*^*th*^ environment, and *ε*_*ijk*_ was the error term. The error term *ε*_*ijk*_ was assumed to follow a normal distribution, 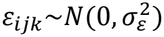, with its associated vector *ε* considered to follow a multivariate normal distribution with covariance matrix 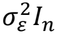. For the rice dataset, *y*_*ijk*_ corresponded to raw phenotypic measurements. LS-means for the genotypes were then computed across all environments using the R package `lsmeans` [61] to estimate a BV per genotype. The Moore-Penrose generalized pseudo-inverse was used during the multiple linear regression procedure to address issues related to aliased variables (i.e., perfect multicollinearity) for certain traits. This adjustment was particularly necessary for traits D, H, and I in the pine dataset.

For the BLUP approaches, the following linear mixed model, was fitted using the R package `lme4` [62]:

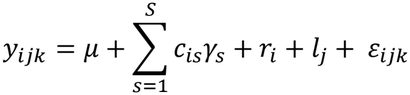

In this model, *y*_*ijk*_, *µ, l*_*j*_ and *ε*_*ijk*_ were defined as in the LS-means approach. Here, *r*_*i*_ represents the random effect of genotype *i* (1 ≤ *i* ≤ *q*), with 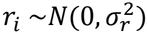, and its associated vector *r* was assumed to follow a multivariate normal distribution with an isotropic covariance matrix, 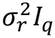. Additionally, to account for population structure, principal component (PC) coordinates for each genotype *i*, denoted *c*_*is*_ (1 ≤ *s* ≤ *S*) were introduced in the model and treated as fixed effects. These PC coordinates were computed using genomic data from each dataset with the R package `mixOmics` [58]. In this context, *γ*_*s*_ represented the effect associated with the *s*^*th*^ PC. Population structure is generally regarded as a low-dimensional process embedded within a high-dimensional space [9, 63]. Consequently, a relatively small number of principal components, rarely exceeding 10, is assumed to effectively capture the underlying genetic structure of the population [10]. The number *S* of principal components (PCs) selected for computing the BLUP values of *r* was chosen to minimize the Akaike information criterion (AIC), obtained via maximum likelihood estimation, within the range of values tested (1 to 10), as AIC is widely regarded as one of the most commonly used criteria for model comparison [64]. The BLUP was then computed using restricted maximum likelihood (REML) to obtain unbiased variance components estimates for use in the BLUP expression. The computed BLUP values using this linear mixed model were named BLUP-PCA values. This model was also fitted without the component 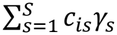, using the REML procedure, to obtain BLUP values which do not account for population structure.

In the WISER framework, the overall mean and environment were modeled as fixed-effect factors for rice. For apple, fixed effects included the mean, environment (country, year, management), and rows and positions, with the latter treated as environment-specific factors for spatial adjustment. For maize and pine, the mean and environment were modeled as fixed factors, while row/column (maize) and GPS coordinates (pine) were treated as quantitative environment-specific variables. This strategy minimized the number of fixed effects and reduced the risk of inestimability. For instance, the maize dataset contained 1,061 environment levels for anthesis, which expanded to 2,122 quantitative variables when including row and column; modeling them as factors would have created an even larger set. Using rows and positions (apple), row/column (maize), and GPS coordinates (pine) directly in the model also avoided the pre-processing step of spatial heterogeneity correction required for LS-means and BLUP. WISER implements three whitening procedures via the R package `whitening` [44]. For each trait, the `optimize_whitening_and_regularization` function automatically selects the optimal whitening method and regularization parameter through a grid search on a data subset, minimizing mean squared error (see Section 2.2).

#### 2.3.3. Compared predictive models

Five predictive models were employed for genomic prediction: random forest (RF), support vector regression (SVR), reproducing kernel Hilbert space (RKHS) regression, and least absolute shrinkage and selection operator (LASSO). These models were selected to represent diverse genetic architectures and to evaluate whether the genomic predictive ability (PA) rankings across the different BV estimation methods and traits remained consistent, regardless of the prediction approach. The RF, SVR, and LASSO models were fitted using the R packages `ranger` [65], `kernlab` [66], and `glmnet` [67], respectively. GBLUP and RKHS regression were fitted using the R package `KRMM` [68]. For RF, the number of randomly selected candidate variables (`mtry`) for testing splits at each tree node was set to one-third of the total number of SNPs, a common choice for regression tasks, and the number of trees was fixed at 1,000. For SVR, LASSO, GBLUP, and RKHS regression, hyperparameters were estimated or selected following the approach described in [68]. Specifically, for SVR, a Gaussian kernel was applied using the `ksvm` function, and its bandwidth parameter was estimated automatically using the heuristic in the `sigest` function. The regularization parameter *C* for SVR was computed as 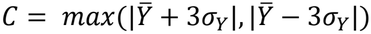, where 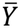 and *σ*_*Y*_ are the mean and standard deviation of the estimated BVs respectively, as recommended by [69]. The *ε*-insensitive loss parameter in SVR was set to its default value of 0.1, as tested values ranging from 0.01 to 0.5 produced similar results, which were comparable to those of other prediction models. For LASSO, the regularization parameter was determined using K-fold cross-validation implemented in the `cv.glmnet` function, with the default value of `nfolds` set to 10. For GBLUP, the `krmm` function was applied with a linear kernel, using the default initial guess values for the variance component estimates required by the EM-REML algorithm. The same settings were used for RKHS regression, except that a Gaussian kernel was applied. The Gaussian kernel’s bandwidth parameter referred to as the rate of decay in `krmm` was set to its default value of 0.1, which provided comparable results to the other prediction models.

A 5-fold cross-validation (CV) scheme with 20 random shufflings of the datasets was employed to estimate the distributions of predictive ability and genomic heritability (*h*^2^) across various prediction models and BV estimation methods for all traits and species. For each shuffle, a new seed was generated by multiplying the shuffle number by 100, thereby introducing substantial variation in the randomized genotype distribution across shuffles. This approach generated a distribution of 100 estimated PA values for each combination of species, trait, prediction model, and BV estimation method. Similarly, a distribution of 100 estimated *h*^2^ values was generated with the GBLUP model using this approach. For each BV estimation method, PA was evaluated for each validation fold by computing the Pearson correlation between the genomically predicted BVs and those estimated using the BV estimation method. In contrast, *h*^2^ was computed from the estimated variance components of the following GBLUP model, which was fitted on the training folds:

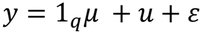

In this model, *y* = (*y*_1_, …, *y*_*q*_)′ represents the vector of *q* BVs estimated from either WISER, LS-means, or BLUP approaches. The vector 1_*q*_ = (1, …,1)^′^ consists of *q* ones, and *µ* corresponds to the fixed overall mean. The vector *u* = ( *u*_1_, …, *u*_*q*_)′ represents the genetic values associated with the *q* genotypes, where 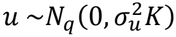, and *K* = *MM*^′^ is the genomic covariance matrix. Here, 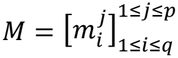 is the centered SNP marker matrix of size *q* x *p*, where 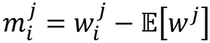, with 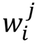 coded as 0, 1, or 2. The error vector *ε* = (*ε*_1_, …, *ε*_*q*_)′ follows a normal distribution,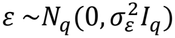. From this model, the genomic heritability *h*^2^ was defined as:

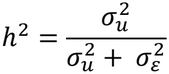

#### 2.3.4. Computed statistics

From the generated distributions of predictive ability (PA) and genomic heritability (*h*^2^), median values were computed for each specific BV estimation method to facilitate graphical representations and subsequent analyses. To explore the relationships between PA and *h*^2^, deviations (i.e., differences) in the PA medians were compared to those in the *h*^2^ medians.

Additionally, for each trait-species combination, deviations in median PA were compared with Cramér’s *V*, computed between genotypes and environments, to assess the relationship between these deviations and the degree of genotype–environment association and the resulting potential confounding between genetic and environmental effects. For each trait-species combination, Cramér’s *V* was defined as:

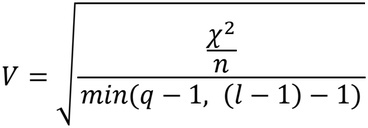

where *χ*^2^ is Pearson’s chi-squared statistic, defined as:

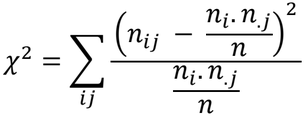

Here, *n* is the total number of observations, *n*_*ij*_ is the number of joint observations for genotype *i* and environment *j, n*_*i*_. = ?_*j*_ *n*_*ij*_ is the number of observations for genotype *i* (*i* = 1, . ., *q*), and *n*_.*j*_ = ?_*i*_ *n*_*ij*_ is the number of observations for environment *j* (*j* = 1, . ., *l* − 1).

Cramér’s *V* provides a measure of the strength of association between genotypes and environments and was therefore used here as an indicator of genotype–environment confounding. It was particularly reported when median PA was low for a given trait-species combination, to investigate whether genotype–environment confounding could contribute to the observed decrease in predictive ability. Cramér’s *V* was computed using the R package `vcd` [70].

In the same manner, the deviations in the PA medians were compared to the Shannon entropy of genotype frequencies across all trait-species combinations. For each trait-species combination, the Shannon entropy of genotype frequencies was defined as:

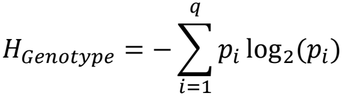

where *p*_*i*_ is the relative frequency of observations of the *i*^*th*^ genotype for the trait in the dataset, and *q* is the total number of genotypes. This comparison was intended to assess the impact of imbalanced genotype frequencies across all trait-species combinations on the deviations observed in the PA medians. A similar comparison was made between these deviations and the Shannon joint entropies of genotype and site, as well as genotype and environment. For each trait-species combination, these joint entropies were computed as follows:

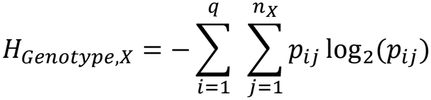

where *X* corresponds to either site or environment, *p*_*ij*_ represents the joint relative frequency of observations of genotype *i* and site (or environment) *j* in the dataset, and *n*_*X*_ is the number of sites (or environments). These joint entropies were also used as complementary indicators of the degree of confounding between genotypes and their respective sites or environments.

For the first scenario of the simulation framework described in section 2.3.1, breeding values (BVs) were estimated using a correctly specified model that included the intercept and all environmental effects as fixed effects. Estimation performance was assessed with and without whitening of the fixed-effect variables using two criteria: estimation accuracy (EA), defined as the Pearson correlation between simulated genetic values and estimated BVs, and mean squared error (MSE), defined as the mean squared difference between simulated genetic values and estimated BVs. In the absence of model misspecification, estimates obtained without whitening are equivalent to classical best linear unbiased estimates (BLUEs) and are therefore referred to as BLUEs throughout this study. For each simulated phenotype dataset, predictive performance was evaluated using both predictive ability (PA) and true predictive ability (TPA). PA was defined as the median of 30 Pearson correlations between estimated BVs and their corresponding GBLUP predictions, obtained from a five-fold cross-validation procedure repeated over six random shufflings of the dataset. TPA was computed analogously, except that correlations were calculated between the simulated genetic values and the corresponding GBLUP predictions trained from either the estimated BVs or the simulated genetic values. Three prediction metrics were therefore considered: (i) PA, (ii) TPA based on GBLUP predictions trained from estimated BVs (WISER or BLUE), and (iii) TPA based on GBLUP predictions trained from the true simulated genetic values (*u*), which represents the upper bound achievable in the absence of estimation error. For the second scenario, only EA and MSE were evaluated.

For the simulated datasets generated under the third scenario and spanning different levels of genotype–environment association, EA and MSE obtained with WISER and BLUE were compared across increasing values of Cramér’s *V*. In addition, the deviation in EA between WISER and BLUE was computed and examined across increasing values of Cramér’s *V* to assess how the benefit of whitening for genetic value estimation changed with increasing genotype–environment confounding.

To explore the relationship between genotype population structure and deviations in PA medians, dimensionality reduction and clustering techniques were employed. UMAP was used to generate five-dimensional representations of the genomic data for each species. The choice of five dimensions was a trade-off between preserving enough genetic variability for effective clustering and mitigating the “curse of dimensionality”, which can reduce the accuracy of distance-based methods in higher-dimensional spaces. The UMAP approach was implemented using the R package `umap` [59]. The resulting five-dimensional representations were clustered using the *K*-means algorithm for each trait-species combination, with the optimal number of clusters determined as the value of *K* that maximized the Calinski-Harabasz (CH) index. Note that the clustering was performed separately for each trait–species combination using only genotypes with available phenotypic measurements, as such data were not consistently available across all genotypes for each trait. Moreover, PCA was not used for dimensionality reduction prior to K-means, as it failed to distinguish known accessions and 27 progeny families in the apple REFPOP dataset (see Supplementary Figures 1 to 4). In contrast, UMAP, without any labeling information, was able to effectively separate the accessions and most of the families, making it better suited for use prior to the clustering algorithm.

The CH index was evaluated over a range of *K* values from 2 to 30 and is defined as:

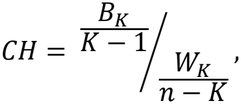

where

- 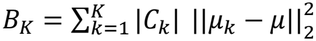 is the between-cluster sum of squares (BCSS) for the SNP marker data. Here, 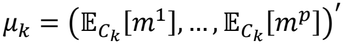 represents the centroid vector for cluster *C, µ* = (E [*m*^1^], …, E[*m*^*p*^])^′^ is the mean vector for the entire dataset, and |*C*_*k*_| is the number of data points (i.e., vectors) in cluster *C*_*k*_.
- 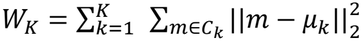 is the within-cluster sum of squares (WCSS), quantifying the compactness of data points around their respective cluster centroids. Here, *m* = (*m*^1^, …, *m*^*p*^)′ represents an SNP vector with values coded as 0, 1 or 2.

The CH index generalizes Fisher’s test statistic from a one-way ANOVA with *K* levels to the multidimensional setting. While this index is not a test statistic and lacks a formal statistical distribution, it serves as a robust metric to balance the compactness of clusters with their separation, providing an effective measure for determining the optimal clustering structure. Note that *K*-means clustering aims to find the optimal partitioning of data into *K* clusters that minimizes *W*_*K*_. Therefore, the CH index is a natural choice for selecting *K*, as both the CH index and *K*-means are related to the total sum of squares (TSS) decomposition, which is expressed as:

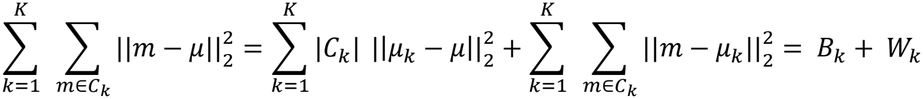

The CH index was computed using the R package `fpc` [71]. Finally, to quantify genetic differentiation between genotypes, the average fixation index *F*_*ST*_ was computed using genomic data associated with the *K* -estimated clusters for each trait-species combination. Notably, for each trait-species combination, the genomic data used accounted for genotype incidence relative to the available phenotypic measures in the experimental design. To improve computational efficiency and minimize resource usage, a stratified sampling approach was employed to maintain the data structure, while limiting the genomic dataset to 5,000 rows per trait-species combination. The *F*_*ST*_ values were calculated using the R packages `adegenet` [72] and `hierfstat` [73].

## 3. Results

### 3.1. Population structure at the genotype level

Supplementary Figures 5 to 16 present a visual assessment of genotype population structure based on the complementary use of heatmaps, PCA and UMAP applied to the SNP data. The apple and pine datasets revealed pronounced population structuring, characterized by distinct clusters corresponding to family groups. Similarly, the maize dataset displayed some clustering patterns despite the absence of prior family information. In contrast, the rice dataset showed minimal clustering, consistent with the findings of [18], who reported no detectable structure in this dataset.

### 3.2. Genomic predictive ability (PA) and heritability (*h*^2^) estimates

Tables 1 and 2 summarize the median PA and *h*^2^ values for BVs estimated using WISER, LS-means, and BLUP approaches across all species and traits, for the GBLUP model. Supplementary File 3 presents median PA values for additional prediction models (random forest, SVR, RKHS, and LASSO), which exhibit, for WISER, LS-means, and BLUP approaches, consistent rankings, similar deviations in median PA values and comparable PA ranges across most models. Therefore, only the GBLUP results are presented here for conciseness. Supplementary Figures 17 to 20 also present a graphical summary of the GBLUP PA values, organized by species, for BVs estimated using WISER, LS-means, and BLUP approaches.

**Table 1.** GBLUP median PA values for BV estimation methods (WISER, LS-means, and BLUP approaches) across all species and traits, with interquartile ranges (IQR) in parentheses. Values are based on 100 cross-validation (CV) scenarios.

| Dataset | Trait | GBLUP median PA values |  |  |  |
|---|---|---|---|---|---|
|  |  | WISER | LS-means | BLUP | BLUP-PCA |
| Rice dataset<br>334 genotypes<br>9,929 SNPs | days to flowering (FL) | 0.74 (0.05) | 0.41 (0.12) | 0.41 (0.11) | 0.21 (0.11) |
|  | plant height (PH) | 0.73 (0.08) | 0.51 (0.09) | 0.51 (0.09) | 0.42 (0.09) |
|  | grain yield (YLD) | 0.62 (0.08) | 0.49 (0.11) | 0.49 (0.11) | 0.42 (0.10) |
|  | grain zinc concentration (ZN) | 0.69 (0.05) | 0.36 (0.16) | 0.36 (0.16) | 0.26 (0.19) |
| Maize dataset<br>246 genotypes<br>41,723 SNPs | Anthesis | 0.74 (0.07) | 0.69 (0.07) | 0.69 (0.07) | 0.49 (0.11) |
|  | anthesis-silking interval | 0.66 (0.07) | 0.37 (0.18) | 0.37 (0.18) | 0.11 (0.20) |
|  | ear height | 0.67 (0.07) | 0.24 (0.15) | 0.24 (0.13) | 0.11 (0.19) |
|  | grain number | 0.68 (0.09) | 0.53 (0.14) | 0.52 (0.14) | 0.34 (0.16) |
|  | grain weight | 0.61 (0.10) | 0.54 (0.13) | 0.54 (0.13) | 0.15 (0.18) |
|  | grain yield | 0.77 (0.08) | 0.66 (0.10) | 0.66 (0.11) | 0.32 (0.16) |
|  | plant height | 0.67 (0.08) | 0.22 (0.15) | 0.22 (0.15) | 0.18 (0.12) |
|  | Silking | 0.70 (0.08) | 0.67 (0.06) | 0.66 (0.07) | 0.40 (0.13) |
|  | tassel height | 0.67 (0.07) | 0.30 (0.14) | 0.30 (0.14) | 0.16 (0.16) |
| Apple dataset<br>534 genotypes<br>50,000 SNPs | color over | 0.78 (0.05) | 0.70 (0.05) | 0.70 (0.05) | 0.48 (0.07) |
|  | flowering begin | 0.92 (0.02) | 0.64 (0.09) | 0.64 (0.09) | 0.55 (0.09) |
|  | flowering end | 0.86 (0.05) | 0.44 (0.10) | 0.44 (0.10) | 0.36 (0.10) |
|  | flowering full | 0.80 (0.04) | 0.40 (0.10) | 0.42 (0.09) | 0.33 (0.09) |
|  | flowering intensity | 0.66 (0.09) | 0.52 (0.09) | 0.52 (0.09) | 0.32 (0.11) |
|  | fruit number | 0.78 (0.04) | 0.67 (0.06) | 0.67 (0.06) | 0.34 (0.10) |
|  | fruit weight | 0.78 (0.04) | 0.65 (0.06) | 0.65 (0.05) | 0.30 (0.10) |
|  | fruit weight single | 0.73 (0.05) | 0.58 (0.07) | 0.58 (0.07) | 0.52 (0.09) |
|  | harvest date | 0.90 (0.03) | 0.76 (0.06) | 0.76 (0.06) | 0.59 (0.07) |
|  | powdery mildew | 0.56 (0.08) | 0.40 (0.10) | 0.40 (0.10) | 0.28 (0.10) |
|  | russet frequency | 0.60 (0.07) | 0.46 (0.09) | 0.46 (0.09) | 0.36 (0.10) |
|  | Scab | 0.56 (0.09) | 0.33 (0.10) | 0.34 (0.09) | 0.21 (0.12) |
|  | trunk diameter | 0.66 (0.06) | 0.50 (0.06) | 0.50 (0.06) | 0.28 (0.09) |
|  | trunk increment | 0.51 (0.07) | 0.49 (0.06) | 0.48 (0.06) | 0.25 (0.12) |
| Pine dataset<br>208 genotypes<br>20,796 SNPs | D | 0.66 (0.10) | 0.08 (0.19) | 0.14 (0.17) | 0.11 (0.19) |
|  | H | 0.72 (0.08) | 0.18 (0.17) | 0.31 (0.16) | 0.23 (0.18) |
|  | I | 0.35 (0.25) | 0.10 (0.21) | 0.31 (0.20) | 0.20 (0.20) |
|  | T4 | 0.39 (0.22) | 0.24 (0.18) | 0.32 (0.20) | 0.30 (0.23) |
|  | T5 | 0.26 (0.25) | 0.14 (0.22) | 0.26 (0.24) | 0.12 (0.24) |
|  | T6 | 0.20 (0.27) | 0.14 (0.24) | 0.20 (0.26) | 0.11 (0.28) |

**Table 2.** GBLUP median *h*^2^ values for BV estimation methods (WISER, LS-means, and BLUP approaches) across all species and traits, with interquartile ranges (IQR) in parentheses. Values are based on 100 cross-validation (CV) scenarios.

| Dataset | Trait | GBLUP median *h*<sup>2</sup> values |  |  |  |
|---|---|---|---|---|---|
|  |  | WISER | LS-means | BLUP | BLUP-PCA |
| Rice dataset<br>334 genotypes<br>9,929 SNPs | days to flowering (FL) | 0.87 (0.02) | 0.53 (0.06) | 0.53 (0.06) | 0.00 (0.00) |
|  | plant height (PH) | 0.86 (0.03) | 0.76 (0.03) | 0.76 (0.03) | 0.70 (0.05) |
|  | grain yield (YLD) | 0.82 (0.04) | 0.73 (0.05) | 0.73 (0.05) | 0.68 (0.06) |
|  | grain zinc concentration (ZN) | 0.84 (0.03) | 0.55 (0.10) | 0.56 (0.10) | 0.42 (0.17) |
| Maize dataset<br>246 genotypes<br>41,723 SNPs | Anthesis | 0.96 (0.02) | 0.93 (0.03) | 0.93 (0.03) | 0.88 (0.04) |
|  | anthesis-silking interval | 0.82 (0.05) | 0.66 (0.11) | 0.66 (0.11) | 0.00 (0.06) |
|  | ear height | 0.69 (0.05) | 0.60 (0.14) | 0.55 (0.15) | 0.00 (0.01) |
|  | grain number | 0.96 (0.02) | 0.86 (0.06) | 0.85 (0.05) | 0.78 (0.09) |
|  | grain weight | 0.65 (0.06) | 0.83 (0.06) | 0.83 (0.05) | 0.00 (0.00) |
|  | grain yield | 0.97 (0.02) | 0.93 (0.04) | 0.92 (0.05) | 0.80 (0.10) |
|  | plant height | 0.68 (0.05) | 0.34 (0.15) | 0.33 (0.15) | 0.00 (0.01) |
|  | Silking | 0.71 (0.04) | 0.92 (0.04) | 0.92 (0.03) | 0.85 (0.06) |
|  | tassel height | 0.68 (0.04) | 0.54 (0.12) | 0.54 (0.12) | 0.00 (0.00) |
| Apple dataset<br>534 genotypes<br>50,000 SNPs | color over | 0.90 (0.03) | 0.84 (0.03) | 0.84 (0.03) | 0.76 (0.05) |
|  | flowering begin | 0.99 (0.00) | 0.94 (0.02) | 0.94 (0.02) | 0.93 (0.03) |
|  | flowering end | 1.00 (0.00) | 0.73 (0.03) | 0.73 (0.03) | 0.69 (0.04) |
|  | flowering full | 0.94 (0.02) | 0.67 (0.06) | 0.69 (0.05) | 0.64 (0.05) |
|  | flowering intensity | 0.77 (0.03) | 0.66 (0.04) | 0.67 (0.04) | 0.56 (0.07) |
|  | fruit number | 0.76 (0.03) | 0.65 (0.04) | 0.65 (0.04) | 0.42 (0.08) |
|  | fruit weight | 0.75 (0.03) | 0.59 (0.03) | 0.59 (0.04) | 0.35 (0.08) |
|  | fruit weight single | 0.93 (0.02) | 0.81 (0.04) | 0.81 (0.04) | 0.79 (0.04) |
|  | harvest date | 0.97 (0.01) | 0.92 (0.02) | 0.92 (0.02) | 0.88 (0.02) |
|  | powdery mildew | 0.74 (0.04) | 0.51 (0.05) | 0.52 (0.05) | 0.36 (0.09) |
|  | russet frequency | 0.81 (0.04) | 0.67 (0.06) | 0.67 (0.05) | 0.54 (0.10) |
|  | Scab | 0.68 (0.04) | 0.44 (0.05) | 0.45 (0.04) | 0.29 (0.06) |
|  | trunk diameter | 0.82 (0.03) | 0.59 (0.04) | 0.59 (0.04) | 0.35 (0.11) |
|  | trunk increment | 0.59 (0.04) | 0.53 (0.04) | 0.53 (0.05) | 0.25 (0.14) |
| Pine dataset<br>208 genotypes<br>20,796 SNPs | D | 0.95 (0.03) | 0.00 (0.17) | 0.03 (0.22) | 0.00 (0.07) |
|  | H | 0.97 (0.01) | 0.30 (0.39) | 0.56 (0.16) | 0.44 (0.52) |
|  | I | 0.73 (0.16) | 0.00 (0.01) | 0.62 (0.22) | 0.00 (0.00) |
|  | T4 | 0.74 (0.14) | 0.54 (0.22) | 0.62 (0.22) | 0.59 (0.24) |
|  | T5 | 0.52 (0.23) | 0.30 (0.44) | 0.48 (0.28) | 0.00 (0.23) |
|  | T6 | 0.28 (0.34) | 0.08 (0.31) | 0.30 (0.35) | 0.00 (0.11) |

Table 1 highlights that WISER consistently achieved the highest GBLUP median PA values across all species and traits. LS-means and BLUP showed similar median PA for most species and traits, except for the Pine dataset, and generally outperformed BLUP-PCA, except in a few isolated cases. BLUP-PCA consistently showed the lowest median PA values in most cases. It is worth noting that, the GBLUP median PA values, for LS-means and BLUP in Table 1, are consistent with the genomic PA values for the corresponding traits reported in [18], [20], [54], and [55], despite differences in CV schemes and associated scenarios in these studies. This emphasizes the improvement that WISER can offer over currently used BVs estimation methods.

For the pine dataset, GBLUP median PA values were notably lower, particularly when BVs were estimated using LS-means and BLUP-PCA. For example, traits D, H, and I showed strong associations between genotypes and environments, as indicated by high Cramér’s *V* values (0.43, 1.00, and 0.41, respectively). This genotype–environment association induced confounding between genetic and environmental effects and, in some cases, severe or perfect multicollinearity among the corresponding dummy variables. For these traits, the variance inflation factor (VIF) for the linear regression underlying LS-means could not be computed, while the restricted maximum likelihood (REML) algorithm used for BLUP-PCA failed to converge to identifiable models. In contrast, WISER mitigated these issues through its whitening transformation, reducing the impact of genotype–environment confounding on genetic value estimation and yielding more stable estimates.

Interestingly, trait H, corresponding to height, is known to be highly heritable. Nevertheless, GBLUP median PA values based on LS-means, BLUP, and BLUP-PCA were low for this trait. A similar pattern was observed for D (duration of budburst), a phenological trait for which genetic variation is also expected [74], for which these three BV estimation methods also yielded lower GBLUP median PA values than WISER. These results suggest that the presence of substantial genetic variation alone does not guarantee accurate downstream genomic prediction when breeding-value estimation is affected by genotype–environment confounding.

In terms of interquartile ranges (IQR) for GBLUP PA values, WISER typically produced smaller ranges compared to LS-means and BLUP, with a few exceptions, most notably in the pine dataset. Across all species and traits, the average IQR values were 0.09 for WISER, 0.12 for LS-means, 0.12 for BLUP, and 0.14 for BLUP-PCA, highlighting WISER’s greater robustness in most scenarios.

To evaluate the potential advantages of WISER over existing BV estimation methods—and given the superior performance of LS-means and BLUP compared to BLUP-PCA—only the differences between the GBLUP median PA values of WISER and those of LS-means and BLUP (referred to as the median PA gain) were calculated for the 33 traits. This yielded 66 median PA gain values. Similarly, differences in GBLUP median *h*^2^ values between WISER and LS-means and BLUP were computed, resulting in 66 median *h*^2^ deviations. The average median PA gain and median *h*^2^ deviation across all species and traits were 0.21 and 0.17, respectively, highlighting WISER’s improved accuracy in BV estimation.

Table 2 shows a similar trend for GBLUP median *h*^2^ values, with WISER producing higher median *h*^2^ values than LS-means and BLUP-based approaches for nearly all species and traits, with the exception of grain weight and silking. However, it is important to note that a higher *h*^2^ estimate does not always indicate a better estimation, as it may result from overfitting of the GBLUP model, which can lead to an overestimation of the trait’s heritability. For instance, while the *h*^2^ values estimated using LS-means and BLUP BVs were higher than those obtained with WISER for grain weight and silking, the corresponding GBLUP median PA values based on LS-means and BLUP BVs were lower than those computed from WISER.

For traits D and I in the pine dataset, both LS-means and BLUP-PCA exhibited exceptionally low *h*^2^ values close to zero due to numerical instability caused by strong confounding between genotypes and environments. Specifically, the estimated values were 0.0035 (0.17) and 0.0042 (0.073) for D, and 0.00092 (0.0086) and 0.0011 (0.0027) for I. When rounded to two decimal places in Table 2, these values appear as zero, but they instead reflect challenges in model convergence rather than a true lack of heritability. For trait D, BLUP also exhibited an exceptionally low *h*^2^ value due to the same challenges. Similarly, convergence difficulties were observed with BLUP-PCA BVs for several other traits across the rice, maize, and pine datasets, where the estimated *h*^2^ was also exceptionally low. In contrast, these issues with both LS-means and BLUP approaches were not observed in the REFPOP dataset, which represented a larger population of 534 genotypes compared to the other datasets: 334 in rice, 246 in maize, and 208 in pine. This larger sample size likely contributed to improved model convergence and more reliable estimation.

Conversely, rounding could also mask very high *h*^2^ estimates. In the apple dataset, WISER yielded a median *h*^2^ of 0.9967 for flowering end, with an interquartile range of only 0.0009 and estimates ranging from 0.9929 (minimum) to 0.9980 (maximum). Consequently, when rounded to two decimal places in Table 2, the median *h*^2^ is reported as 1.00 and the IQR as 0.00. Thus, the value of 0.00 reported for the IQR reflects the very limited variability among estimates, rather than an actual zero and absence of variability.

It is also worth noting that flowering end and flowering full, like flowering begin, are highly heritable traits. However, flowering end and flowering full were phenotyped exclusively in Spain, under two management conditions (1 and 2), with flowering end evaluated over four years (2019–2021 and 2023) and flowering full over three years (2019–2021). This more restricted environmental representation may increase the potential for associations between genotype distribution and environmental conditions. Consistent with this possibility, flowering end and flowering full showed Cramér’s *V* values of 0.26 and 0.33, respectively, among the highest values observed across traits in the apple dataset, together with scab (0.31). Although these values do not indicate particularly strong associations in absolute terms, their relative magnitude within the apple dataset suggests greater genotype– environment association for these traits compared with most other traits evaluated. The combination of high trait heritability and comparatively greater genotype–environment association could therefore contribute to the higher median *h*^2^ and PA estimates obtained with WISER relative to LS-means, BLUP, and BLUP-PCA. More generally, the range of Cramér’s *V* values observed across traits differed substantially among species, spanning [0.27, 0.28], [0.23, 0.24], [0.10, 0.33], and [0.41, 1.00] for rice, maize, apple, and pine, respectively. These differences are consistent with the broader pattern observed throughout the study, whereby the relative advantage of WISER tended to increase under stronger genotype–environment association and confounding.

### 3.3. Factors contributing to the variations in median PA gain

To further investigate the factors contributing to the variations in median PA gain across species and traits, correlations were computed between the median PA gain and the median *h*^2^ differences (i.e., deviations), as well as with several additional statistics defined in Section 2.3: Cramér’s *V* between genotypes and environments, entropy of genotype frequencies in datasets, Shannon joint entropy of genotype and site (or environment) in datasets, and the *F*_*ST*_ calculated from the clusters estimated through the combination of UMAP and *K*-means applied to SNP marker data. These analyses provide insights into the factors associated with the observed differences between WISER and the alternative methods.

Figure 1 presents correlation plots between the 66 median PA gain values and various statistics, as defined in Section 2.3.4, for the 33 traits-species combinations. To enable correlation with the 66 PA gain values, each of the 33 statistic values was duplicated.

**Figure 1.**
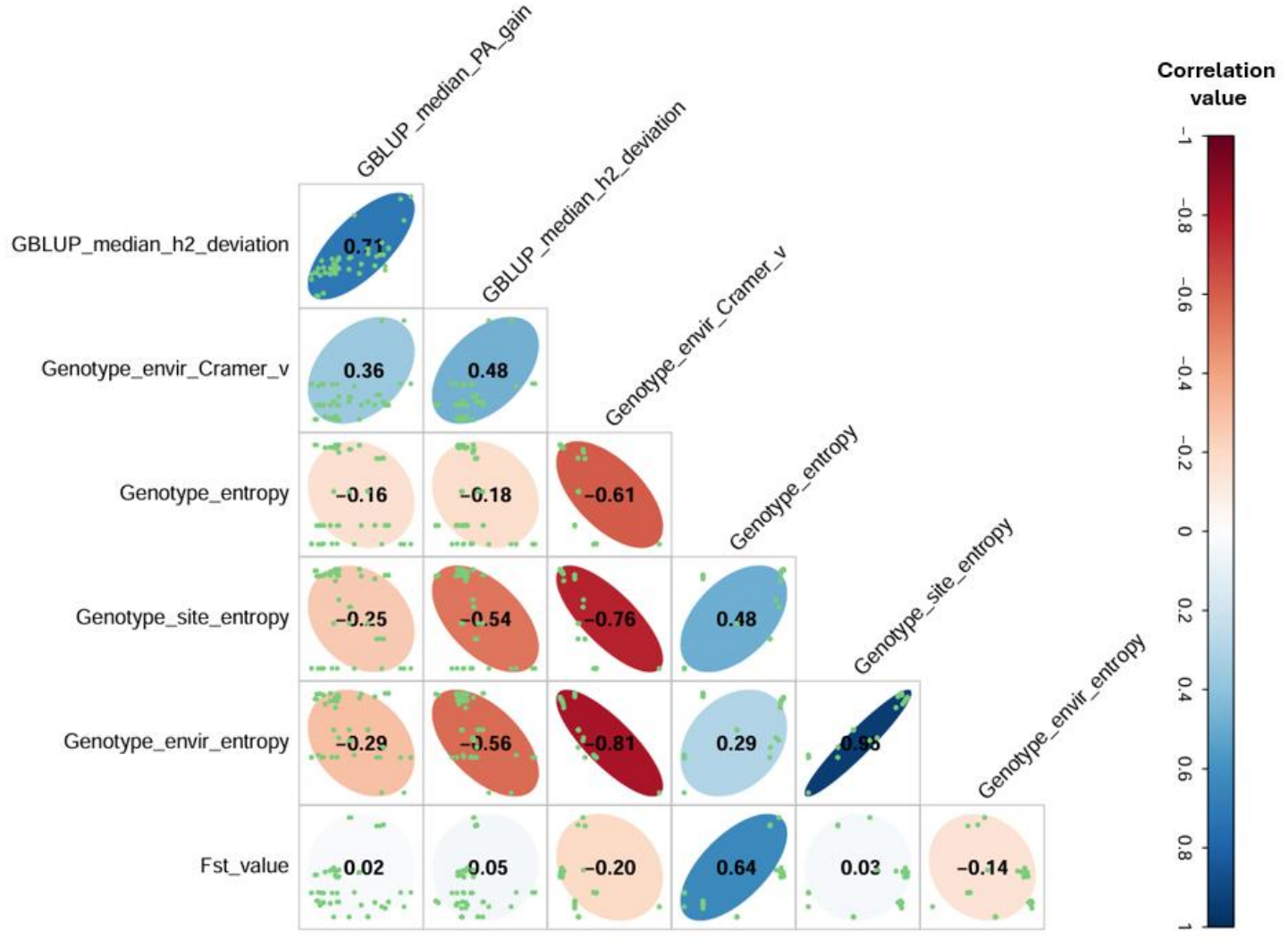
Correlation plots between GBLUP median PA gain, median *h*^2^ deviations, Cramér’s *V*, Shannon entropies, and *F*_*ST*_ values across all species and traits.

A strong positive correlation of 0.71 was observed between the median PA gain values and the median *h*^2^ deviations. This result aligns with expectations, as an increase in *h*^2^ estimates typically reflects higher genetic variance estimates and better predictive ability. In contrast, a weak-to-moderate positive correlation (0.36) was found between the median PA gain values and Cramér’s *V* between genotype and environment. Similarly, weak negative correlations were found between the median PA gain values and the Shannon entropy of genotype frequencies across datasets (-0.16), as well as the Shannon joint entropies of genotype and site (-0.25) and genotype and environment (-0.29). Although these correlations were relatively weak, they suggested a trend whereby the median PA gain achieved by WISER tended to increase in scenarios with greater genotype-site or genotype–environment confounding. Notably, when considering the median *h*^2^ deviations rather than the median PA gain, stronger associations emerged. A moderate positive correlation (0.48) was found between the median *h*^2^ deviations and Cramér’s *V* between genotype and environment, while moderate negative correlations with the Shannon joint entropies of genotype and site (-0.54) and genotype and environment (-0.56) were observed, suggesting a greater impact of genotype-site and genotype– environment confounding on genetic variance estimation.

On the other hand, correlations close to zero were observed between the median PA gain and *F*_*ST*_, as well as between the median *h*^2^ deviations and *F*_*ST*_, making the relationship inconclusive. The weak negative or near-zero correlations observed in Figure 1 are likely unreliable and may reflect species-specific factors such as marker density, linkage disequilibrium patterns, and allele or genotype frequencies, which show little to no variation across traits within a given species. Table 3 presents the median values and interquartile ranges (IQR) of Shannon entropy for genotype frequencies (*H*_*Genotype*_) and *F*_*ST*_ values across traits for each species. Notably, it shows limited or no variation in these statistics among traits within each species, which likely contributed to the weak negative or near-zero correlations.

**Table 3.** Median and interquartile range (IQR) values for Shannon entropy of genotype frequencies (*H*_*Genotype*_) and average fixation index (*F*_*ST*_) across traits per species.

| Species | $median(H_{Genotype})$ | $IQR(H_{Genotype})$ | $median(F_{ST})$ | $IQR(F_{ST})$ |
|---|---|---|---|---|
| Rice | 8.365 | 0.000 | 0.010 | 0.000 |
| Maize | 7.932 | 0.001 | 0.025 | 0.003 |
| Apple | 8.929 | 0.042 | 0.056 | 0.006 |
| Pine | 7.695 | 0.005 | 0.035 | 0.001 |

Figure 2 presents correlation plots for the apple dataset, generated using the same approach as in Figure 1. This dataset included the highest number of traits (14) among all species. The plots are based on 28 values for the median PA gain and median *h*^2^ deviation, calculated from the differences between median values of WISER and BLUP, as well as WISER and LS-means. This enabled an exploration of intra-species relationships within a single species. The observed trends were consistent with those from the inter-species analysis, particularly regarding correlations between median PA gain, median *h*^2^ deviation, Cramér’s *V*, and the Shannon entropies. A positive correlation between median PA gain and *F*_*ST*_ was also observed, however, this may be coincidental given the limited variability and dispersion seen in the corresponding plot.

**Figure 2.**
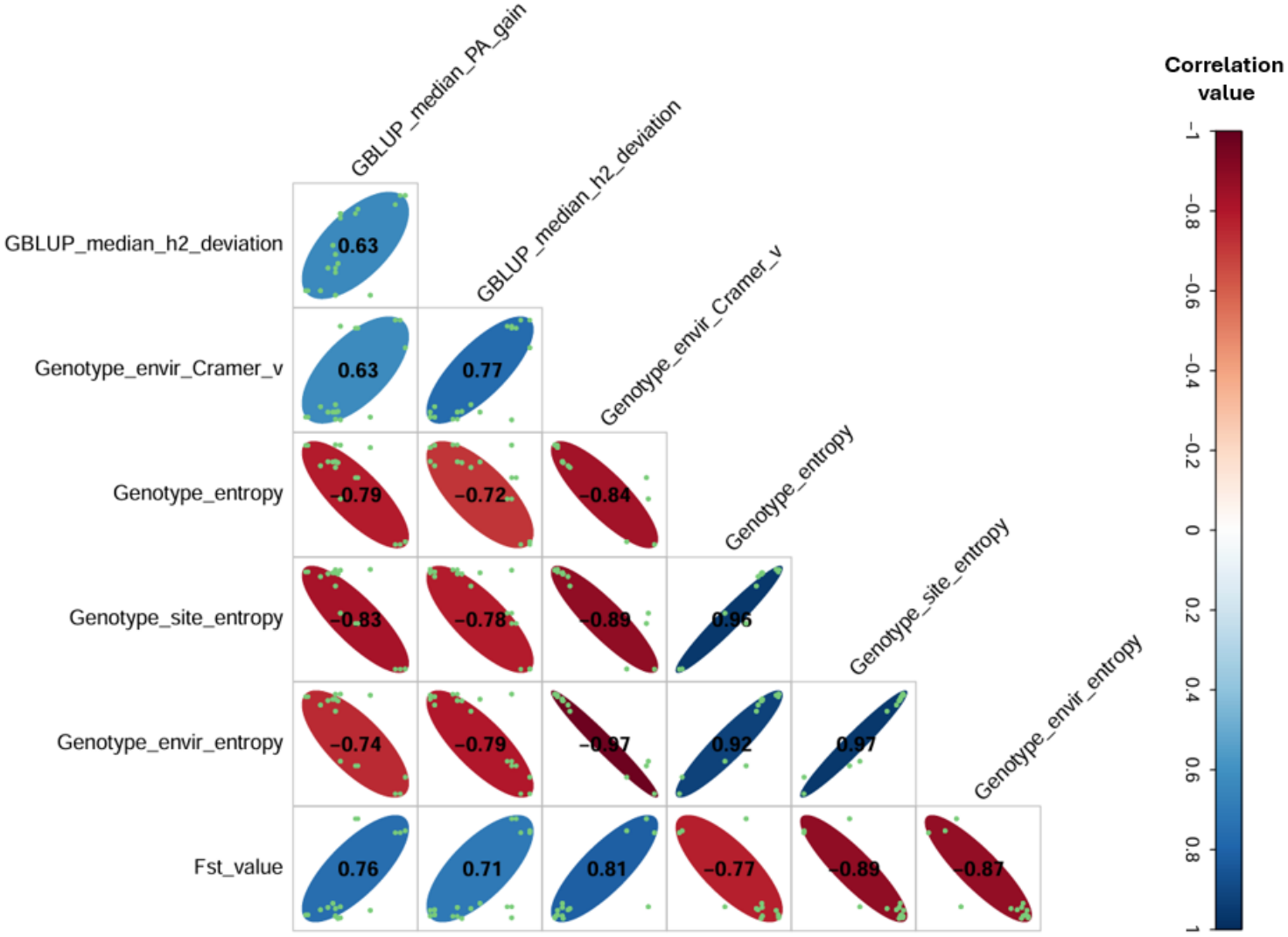
Correlation plots between GBLUP median PA gain, median *h*^2^ deviations, Cramér’s *V*, Shannon entropies, and *F*_*ST*_ values for apple traits.

In summary, the positive and negative correlations observed between the median PA gain values (or median *h*^2^ deviations) and Cramér’s *V* and Shannon entropies, respectively, suggest that the benefits of WISER for BV estimation become more pronounced in scenarios characterized by stronger genotype-site or genotype-environment confounding.

### 3.4. Impact of whitening versus non-whitening of fixed-effect variables on genetic value estimation and prediction

Table 4 presents the median and interquartile range (IQR, shown in parentheses) of estimation accuracy (EA), mean squared error (MSE), predictive ability (PA), and true predictive ability (TPA) across 100 simulated datasets for the trait with moderate heritability (0.5) under the first simulation scenario described in section 2.3.1. In this scenario, breeding values were estimated using a correctly specified model that included the intercept and all environmental effects as fixed effects, with and without whitening of the fixed-effect variables (WISER and BLUE, respectively). In Table 4, the column *u* corresponds to GBLUP predictions trained from the true simulated genetic values. Consequently, EA MSE, and PA are not defined for this reference and are therefore not reported.

**Table 4.** Median and IQR of EA, MSE, PA, and TPA across 100 simulated datasets for the trait with moderate heritability (0.5), by species, under WISER (whitening) and BLUE (no whitening). TPA is also reported for GBLUP predictions trained using the true simulated genetic values (*u*).

| Dataset | Statistic | WISER | BLUE | *u* |
|---|---|---|---|---|
| Rice | EA | 0.82 (0.04) | 0.82 (0.03) | – |
|  | MSE | 0.22 (0.02) | 0.23 (0.03) | – |
|  | PA | 0.57 (0.08) | 0.57 (0.07) | – |
|  | TPA | 0.69 (0.06) | 0.70 (0.06) | 0.80 (0.05) |
| Maize | EA | 0.65 (0.05) | 0.40 (0.13) | – |
|  | MSE | 0.29 (0.06) | 2.43 (1.52) | – |
|  | PA | 0.37 (0.12) | 0.16 (0.12) | – |
|  | TPA | 0.48 (0.15) | 0.36 (0.19) | 0.62 (0.1) |
| Apple | EA | 0.77 (0.04) | 0.77 (0.04) | – |
|  | MSE | 0.31 (0.03) | 0.34 (0.03) | – |
|  | PA | 0.50 (0.09) | 0.50 (0.09) | – |
|  | TPA | 0.65 (0.08) | 0.65 (0.08) | 0.76 (0.06) |
| Pine | EA | 0.89 (0.03) | 0.26 (0.12) | – |
|  | MSE | 0.12 (0.02) | 7.03 (5.24) | – |
|  | PA | 0.38 (0.10) | -0.21 (0.14) | – |
|  | TPA | 0.43 (0.11) | 0.18 (0.15) | 0.45 (0.10) |

The results indicate that whitening generally improved genetic value estimation. Median EA values were higher for WISER than for BLUE in the maize and pine datasets and were similar between the two methods in the rice and apple datasets. Consistently, median MSE values were lower for WISER than for BLUE in the maize and pine datasets and comparable between methods in the rice and apple datasets. Similar trends were observed for prediction, with both PA and TPA showing higher median values for WISER than for BLUE in the maize and pine datasets and little difference between methods in the rice and apple datasets. In all cases, PA based on WISER-estimated BVs remained below the corresponding TPA, which in turn remained below the TPA obtained from GBLUP predictions trained on the true simulated genetic values (*u*). This latter metric represents the upper bound of predictive performance achievable in the absence of estimation error. Overall, these results are consistent with those observed in the real datasets and further demonstrate that whitening fixed-effect variables can improve both genetic value estimation and subsequent prediction when fixed effects are genetically confounded. Supplementary Figures 21 to 24 provide graphical summaries of these results, organized by species.

Under the second scenario, in which only the intercept was included as a fixed effect, resulting in a naïve estimation of *u*, EA medians were low and identical between WISER and BLUE across all species (Table 5). MSE medians were also high and nearly identical between the two methods. Thus, under this misspecified model, in which environmental effects were omitted, WISER and BLUE produced nearly identical estimates, indicating that whitening the intercept alone provided no improvement in genetic value estimation. Supplementary Figures 25 to 28 provide graphical summaries of these results, organized by species.

**Table 5.**
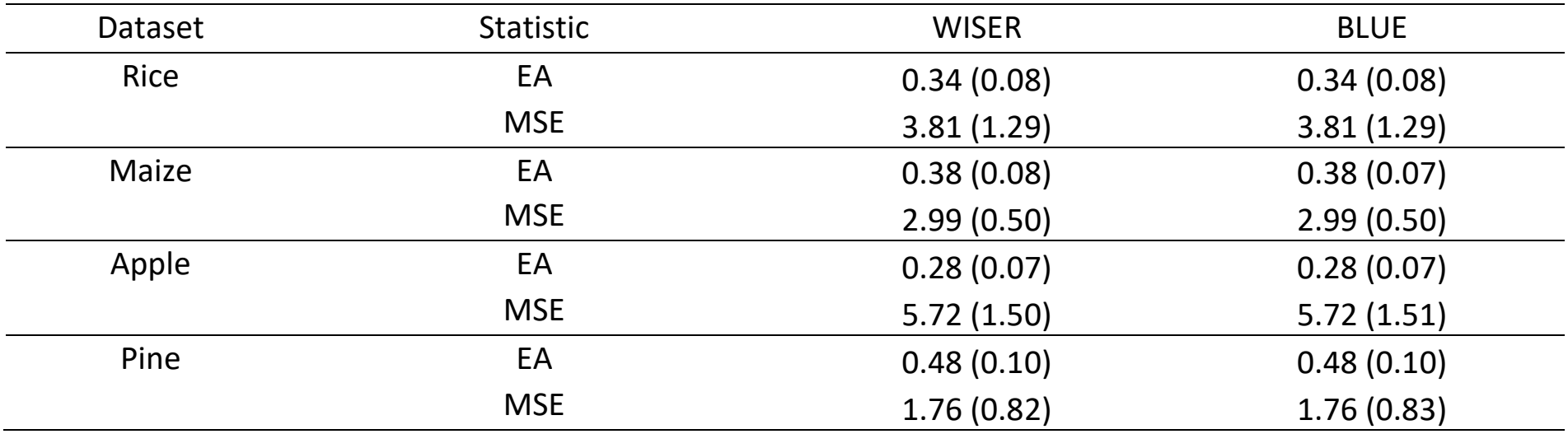
Median and interquartile range (IQR) of estimation accuracy (EA) and mean squared error (MSE) for simulated genetic values by species under WISER (whitening) and BLUE (without whitening) in the second simulation scenario.

| Dataset | Statistic | WISER | BLUE |
|---|---|---|---|
| Rice | EA | 0.34 (0.08) | 0.34 (0.08) |
|  | MSE | 3.81 (1.29) | 3.81 (1.29) |
| Maize | EA | 0.38 (0.08) | 0.38 (0.07) |
|  | MSE | 2.99 (0.50) | 2.99 (0.50) |
| Apple | EA | 0.28 (0.07) | 0.28 (0.07) |
|  | MSE | 5.72 (1.50) | 5.72 (1.51) |
| Pine | EA | 0.48 (0.10) | 0.48 (0.10) |
|  | MSE | 1.76 (0.82) | 1.76 (0.83) |

Under the third scenario, EA decreased and MSE increased for both WISER and BLUE as Cramér’s *V* between genotypes and environments increased. Although both approaches exhibited similar overall trends, the decline in EA was more pronounced for BLUE when Cramér’s *V* exceeded 0.8. Likewise, BLUE showed a sharper increase in MSE at high levels of genotype-environment association. As a result, the EA gain, defined as the difference between the EAs obtained with WISER and BLUE, increased with increasing Cramér’s *V*. Positive correlations of 0.63, 0.41, 0.66, and 0.54 were observed between EA gain and Cramér’s *V* for rice, maize, apple, and pine, respectively. These results suggest that the relative advantage of WISER over BLUE increased as genotype-environment confounding became stronger, particularly at high levels of association.

Figures 3-5 provide graphical representations of these results for the four species. It should be noted that all analyses were performed with heritability fixed at 0.5. Consequently, the reported patterns should not be interpreted as encompassing the full range of EA and MSE behaviors that may arise under different heritability levels as genotype-environment confounding increases.

**Figure 3.**
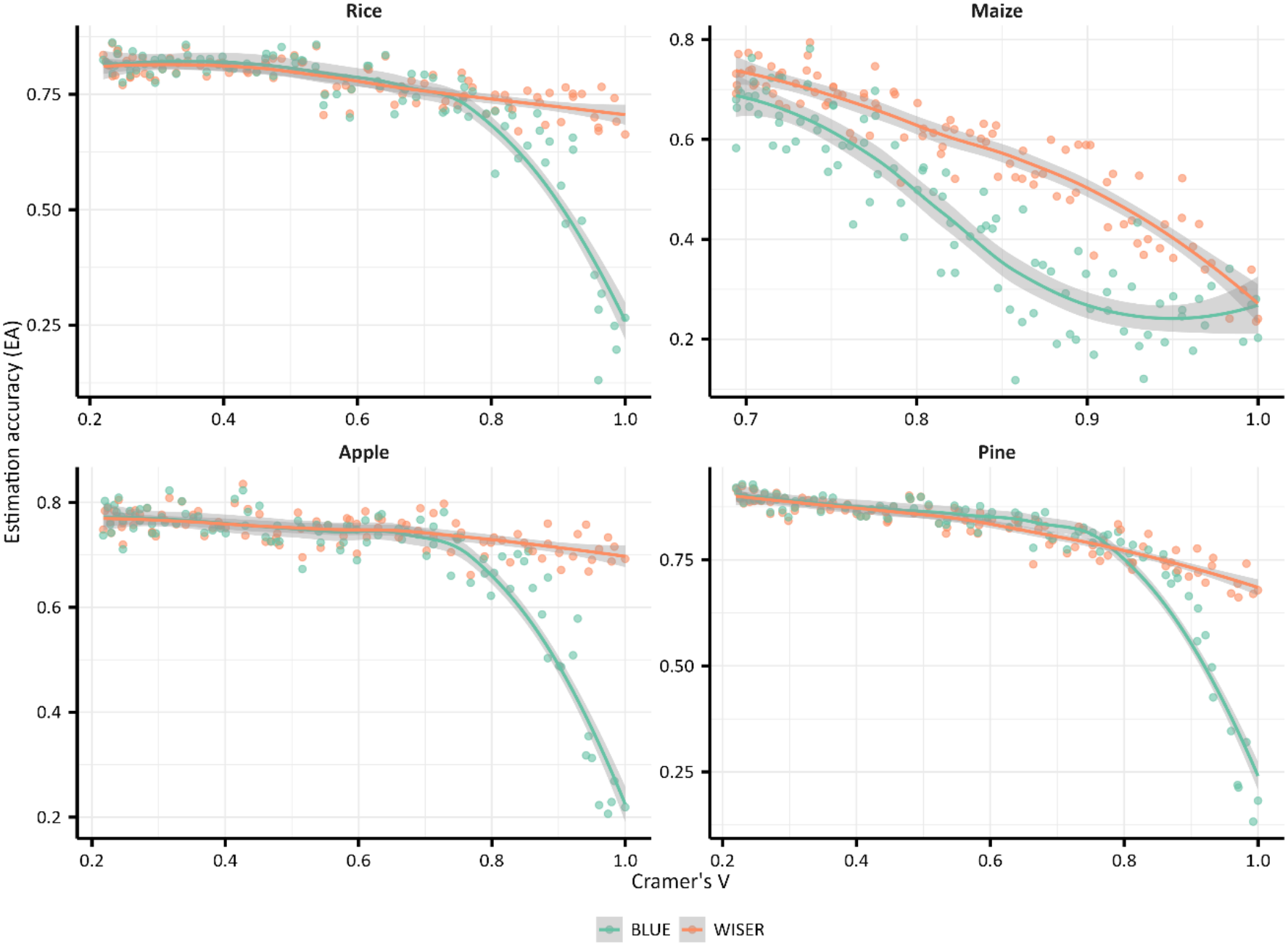
Estimation accuracy (EA) between simulated and estimated genetic values for WISER and BLUE across species, for a simulated trait based on genomic and experimental design data (*h*^2^ = 0.5), at different levels of genotype–environment association (i.e., confounding).

**Figure 4.**
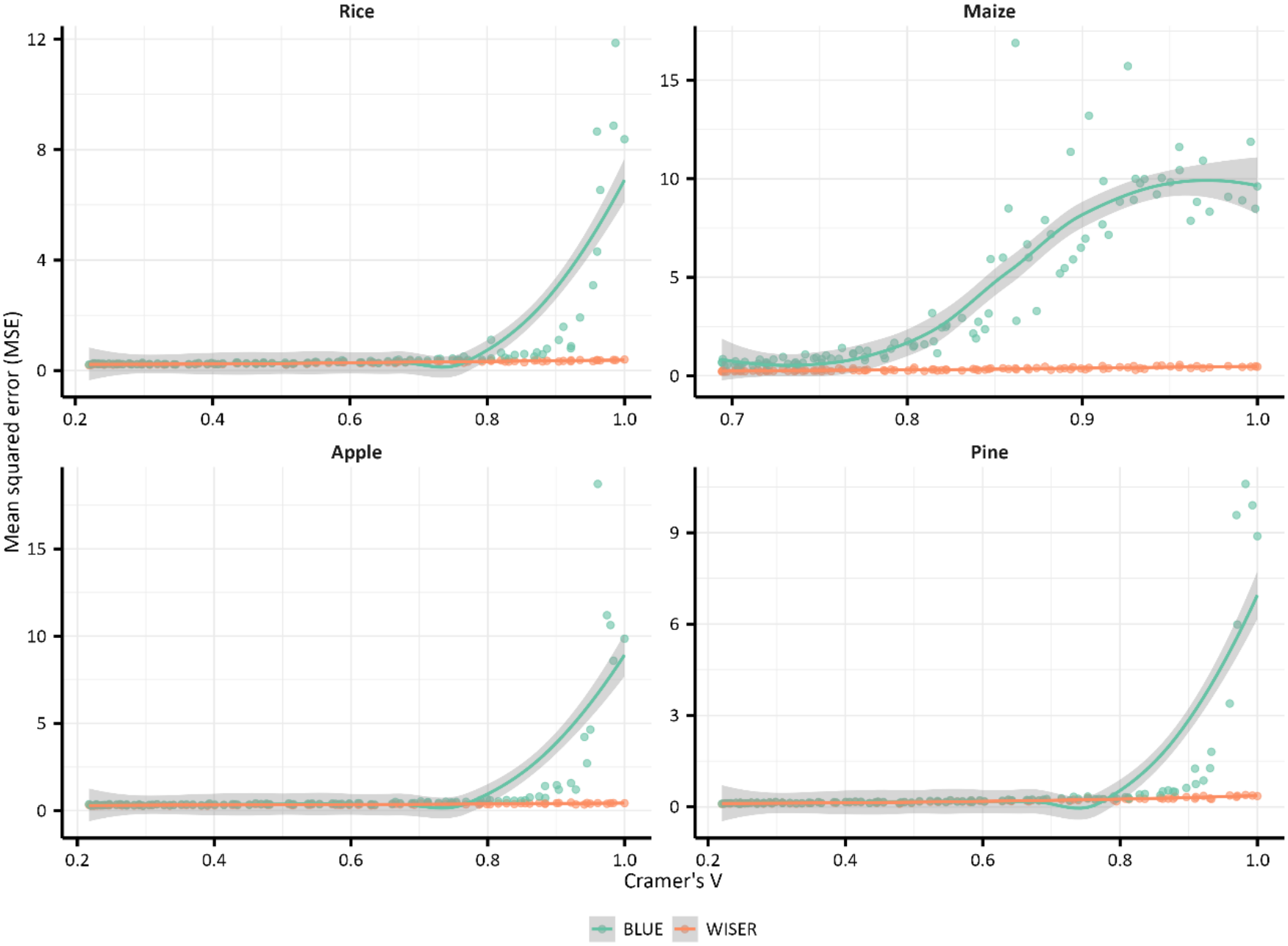
Mean square error (MSE) between simulated and estimated genetic values for WISER and BLUE across species, for a simulated trait based on genomic and experimental design data (*h*^2^ = 0.5), at different levels of genotype–environment association (i.e., confounding).

**Figure 5.**
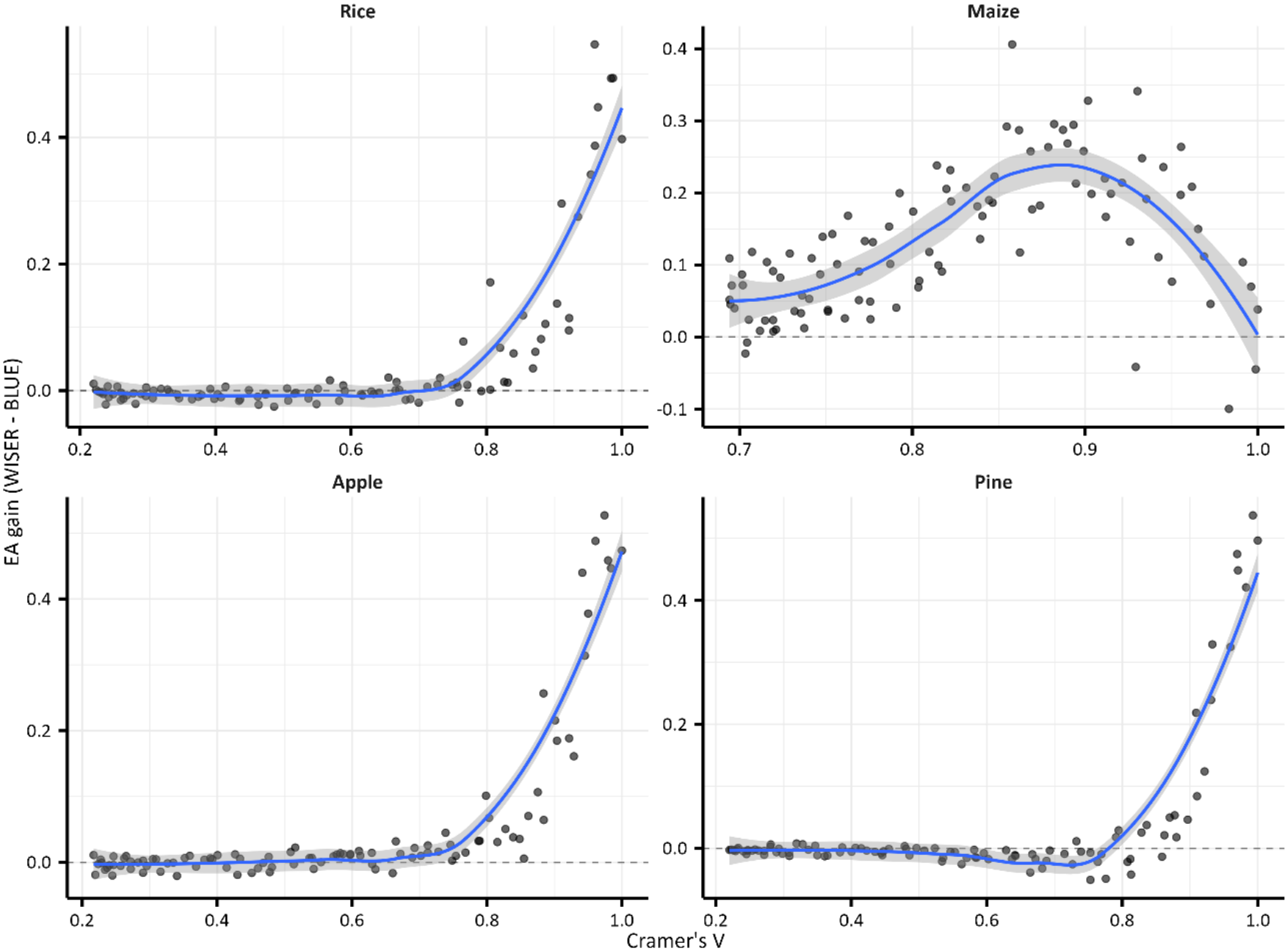
EA gain (EA gain = EA[WISER] - EA[BLUE]) between simulated and estimated genetic values across species, for a simulated trait based on genomic and experimental design data (*h*^2^ = 0.5), at different levels of genotype–environment association (i.e., confounding).

## 4. Discussion

This study provides evidence that WISER can improve BV estimation and downstream prediction relative to conventional approaches such as LS-means, BLUP, and BLUE. These improvements were observed across four datasets encompassing 33 traits and were further supported by simulations representing different levels of genotype–environment confounding derived from the four datasets. Overall, the results indicate that WISER can improve BV estimation and prediction under genotype– environment confounding and may also offer advantages in datasets characterized by imbalanced genotype representation or complex population structure.

### 4.1. Comparison of WISER, LS-means, BLUP, and BLUE

Across the four species and 33 traits considered in this study, WISER consistently yielded higher median predictive abilities (PA) and genomic heritabilities (*h*^2^) than LS-means and BLUP-based approaches. WISER achieved the highest median GBLUP PA for all species, whereas BLUP-PCA frequently exhibited the lowest predictive performance. Taken together, these results indicate that explicitly accounting for the genetic covariance structure during BV estimation can substantially improve downstream genomic prediction.

The simulation study provides further support for this interpretation. Because the true genetic values were known, the estimation accuracy (EA) and true predictive ability (TPA) could be evaluated directly. Across simulation scenarios, whitening was generally associated with higher EA and TPA and lower mean squared error (MSE), although this advantage was not observed in every simulation setting. The benefit of whitening for EA became more pronounced as genotype–environment confounding increased. Importantly, the covariance structure used for whitening was itself estimated under non-optimal conditions and therefore provided only an imperfect representation of the underlying genetic structure. Despite this limitation, whitening generally improved BV estimation under substantial genotype–environment confounding. Although the magnitude of the improvement varied among species and simulation settings, these results suggest that whitening can improve the separation of genetic and environmental sources of variation when confounding is substantial, even when the covariance structure used for whitening is imperfectly estimated. Importantly, these simulations indicate that the advantages of WISER are not limited to downstream predictive performance but can also be associated with more accurate estimation of the underlying genetic values.

Beyond its ability to reduce genotype–environment confounding by accounting for the genetic covariance structure, several other features of WISER may contribute to these improvements. Unlike BLUP-based approaches, WISER does not model the BV vector as a random effect with an imposed covariance structure. Instead, BVs are obtained through a successive OLS procedure after whitening the fixed-effect variables. Although genomic information enters the estimation through the covariance matrix used for whitening, the resulting BVs are not parameterized as functions of individual marker effects. WISER therefore does not impose shrinkage directly on the BVs during their estimation. This distinction may be particularly relevant in unbalanced datasets, where BLUP estimates for poorly represented genotypes can undergo stronger shrinkage toward the population mean. As emphasized by [75], such behavior can alter genotype rankings and may have consequences when estimated BVs are subsequently used as response variables. All experimental designs analyzed here exhibited some degree of incompleteness or imbalance (Supplementary File 4), suggesting that this mechanism could have contributed to the observed differences between methods.

LS-means present a different limitation in the context of BV estimation. By estimating marginal means under an equal-weighting scheme across the levels of other model factors, they may differ substantially from the empirical genotype means when the experimental design is unbalanced.

Although this adjustment is appropriate when the objective is to compare genotypes under common experimental conditions, the resulting LS-means may differ substantially from genotype means derived from their actual representation across environments, with potential consequences for subsequent genomic prediction [76]. WISER instead addresses confounding through whitening of the fixed-effect design while retaining an OLS-based estimation framework.

The comparison with BLUP-PCA also provides insight into the mechanism underlying WISER. Incorporating principal component coordinates as fixed-effect covariates did not improve BV estimation and generally resulted in lower predictive abilities than BLUP or LS-means. This observation is consistent with the theoretical developments presented in Supplementary File 1. Whereas PC-based adjustment relies on a selected subset of eigendirections, WISER uses the full genetic covariance structure through whitening. Rather than explicitly removing or adjusting for only a predefined number of principal components, WISER continuously reweights the eigenspace according to the spectral structure of the covariance matrix. The increasing advantage of WISER with increasing Cramér’s *V* is consistent with this interpretation, as stronger genotype-environment confounding may be more strongly reflected in dominant directions of genetic variation. This removes the need to choose an arbitrary number of PCs and may provide a more comprehensive adjustment for population structure.

Several complementary analyses suggest that the gains observed with WISER are unlikely to result solely from a repeated use of genomic information. First, a sensitivity analysis on the rice dataset (Supplementary File 5, Table S5) showed that WISER retained a substantial part of its advantage when only 100 randomly sampled SNPs were used to estimate the covariance matrix for whitening, while all 9,929 markers remained available for genomic prediction. Despite this strong reduction in the information used to characterize population structure, WISER still yielded higher predictive abilities than LS-means and BLUP for several traits. At the same time, the lower performance observed for grain yield in this sensitivity analysis suggests that the effectiveness of whitening depends on the ability of the marker subset to adequately capture the underlying genetic structure. An incomplete characterization of population structure may therefore reduce the effectiveness of the whitening correction.

Second, an additional two-stage GBLUP analysis (Supplementary File 6) showed that repeatedly incorporating genomic covariance information does not, by itself, reproduce the gains obtained with WISER. Across both the rice and Scots pine datasets, predictive abilities and heritability estimates obtained from the second-stage GBLUP remained nearly identical to those obtained from the original BLUP estimates. The theoretical developments presented in Supplementary File 6 further show that the predictive performance of the second-stage GBLUP remains fundamentally constrained by the quality of the BVs estimated during the first stage. Consequently, reintroducing genomic covariance information in a second prediction step does not inherently generate additional predictive signal or substantially improve heritability estimates.

Taken together, the simulation results, the 100-SNP sensitivity analysis, and the two-stage GBLUP comparison support the hypothesis that a substantial part of the advantage of WISER arises from the whitening-based adjustment of fixed effects and the associated reduction of genotype-environment confounding, rather than merely from a repeated use of genomic covariance information.

Finally, despite relying only on one-dimensional spatial covariates (e.g., row–column positions or GPS coordinates) modeled as fixed effects, WISER consistently yielded higher predictive abilities than LS-means and BLUP approaches that incorporated more sophisticated spatial corrections. In contrast, the other BV estimation methods incorporated both one- and two-dimensional components in their spatial correction. A potential improvement for WISER would be to adopt the approach proposed by [40], which involves modeling spatial heterogeneity as a random effect. This allows for the efficient inclusion of two-dimensional spatial components, which would be impractical to model as fixed effects due to the high number of levels. Furthermore, [40] demonstrated that fitting all relevant effects within a single model generally improves the estimation of genetic values.

One limitation of the WISER framework is the reduced interpretability of fixed-effect estimates after whitening. Although this has little consequence when the primary objective is BV estimation, it may be restrictive in situations where environmental effects themselves are of direct scientific or operational interest.

### 4.2. Impact of confounding effects and imbalanced frequencies

This study also investigated how confounding and imbalances in genotype representation across environments affect BV estimation. The results indicate that WISER is particularly effective under such conditions. For example, in the Scots pine dataset, traits such as D, H, and I exhibited strong genotype– environment associations and confounding, and WISER produced BV estimates associated with higher predictive abilities than LS-means and BLUP-PCA. In these cases, the competing methods were affected by numerical and design-related difficulties, including convergence problems for REML in BLUP-PCA and multicollinearity in the linear models used to derive LS-means.

These observations are important because genotype–environment confounding, particularly when accompanied by pronounced differences in genotype frequencies across environments, can compromise the separation of genetic and environmental effects. Moreover, unequal representation of genotypes across environments is intrinsically related to the covariance structure of the experimental design. Confounding, imbalance, and genetic structure should therefore not necessarily be viewed as independent features, but rather as interacting characteristics that collectively influence BV estimation. Through whitening, WISER modifies the geometry of the fixed-effect design relative to the genetic covariance structure, which may help reduce the impact of dependencies between genetic and environmental effects on BV estimation. This interpretation is consistent with the observed improvements in BV estimation under increasing genotype-environment confounding.

Importantly, some degree of genotype–environment association, imbalance in genotype representation, and population structure is difficult to avoid in real breeding datasets, even when experimental designs are carefully constructed. This is illustrated by the apple REFPOP dataset, which, despite being designed as a reference population for apple breeding, still exhibited measurable levels of confounding and imbalance. This observation highlights the practical relevance of methods capable of accommodating such departures from idealized balanced designs.

### 4.3. Impact of the degree of population structure among genotypes

The study further explored whether the improvement in BV estimation achieved with WISER was related to the degree of population structure among genotypes, quantified through measures of genetic differentiation. No clear general relationship emerged, partly because population structure among genotypes is species-specific and genomic structure varied little, or not at all, among traits within a given species.

Nevertheless, the results suggest that the benefit of WISER is more closely related to the joint distribution of genotypes across environments, and hence to the strength of genotype–environment associations, than to population structure among genotypes considered in isolation. Among the statistics investigated, the strongest correlations with median PA gain were observed for the Shannon joint entropy of genotype and environment and Cramér’s *V*. The correlation with joint entropy was negative, with lower values indicating a less diverse distribution of genotype–environment combinations and, consequently, more restricted genotype representation across environments. Conversely, the correlation with Cramér’s *V* was positive, with increasing values indicating stronger genotype–environment associations.

These results emphasize the importance of considering not only population structure among genotypes but also how genotypes are represented across experimental environments. Population structure may affect genomic prediction when genetic structure is associated with environmental structure, and it is also a well-established source of confounding in association studies [6–12]. WISER provides a framework for accounting for genetic covariance when adjusting for environmental effects during BV estimation.

### 4.4. BV estimation, genetic variance, and predictive ability

The relationship between median PA gain and median *h*^2^ deviation further illustrates the connection between BV estimation, estimated genetic variance, and downstream predictive performance. Across species and traits, a strong positive correlation (0.71) was observed between these two quantities. Thus, cases in which WISER produced larger increases in genomic heritability also tended to be those in which it produced larger improvements in predictive ability.

This association is consistent with the hypothesis that improved separation of genetic and non-genetic sources of variation during BV estimation can increase the genetic signal available for downstream genomic prediction. However, because genomic heritability and predictive ability are themselves related quantities, this correlation should not be interpreted as direct evidence of causality. The simulation results provide additional support for this interpretation. The concurrent increase in estimation accuracy (EA) and decrease in mean squared error (MSE) observed under whitening suggest that the higher genomic heritabilities obtained with WISER do not simply reflect inflation of the genetic variance component, but are often accompanied by more accurate estimates of the underlying genetic values.

Median *h*^2^ deviations were also positively correlated with Cramér’s *V* and moderately negatively correlated with the Shannon joint entropy of genotype and site or environment. Similar, although generally weaker, trends were observed for median PA gain. These relationships were particularly pronounced in the apple dataset. Together with the simulation results, these findings suggest that WISER provides its greatest benefits when genotype distribution is strongly dependent on site or environment, conditions under which separating genetic and environmental sources of variation becomes particularly challenging.

### 4.5. Potential for genome-wide association studies (GWAS)

Although GWAS applications were not formally evaluated in this study, the WISER framework may also have implications for genome-wide association studies. Preliminary analyses (results not shown) indicated that several known QTLs in apple were detected when either LS-means or WISER-estimated BVs were used as response variables, while additional QTLs differed between the two BV estimation approaches.

These preliminary observations suggest that comparing association signals obtained from WISER-estimated BVs with those obtained using conventional phenotype-adjustment methods could provide useful complementary evidence. However, the detection of additional associations should not by itself be interpreted as evidence that these associations are true QTLs or that WISER necessarily improves GWAS performance. A dedicated evaluation of false-positive rates, statistical power, effect-size estimation, and QTL replication would be required to establish this.

Accordingly, further investigation of WISER for association mapping represents an important direction for future research but is beyond the scope of the present study.

### 4.6. Limitations, computational efficiency, and population size considerations

The advantages of WISER are expected to be greatest when several fixed-effect variables contribute to the definition of the environmental structure. Its potential benefit is more limited when the fixed-effect design contains little information to whiten. This was illustrated by the second simulation scenario, in which the intercept was the only fixed effect. Under this configuration, WISER and BLUE produced nearly identical estimates of *u*, indicating that whitening provides no additional information when the fixed-effect space is essentially restricted to the intercept.

This result clarifies an important aspect of the method: WISER does not inherently improve BV estimation in every experimental design. Its benefit arises when the fixed-effect design contains environmental or experimental structure that is non-trivially related to the genetic covariance structure. Consequently, richer and appropriately specified experimental designs provide greater opportunity for whitening to improve the separation of genetic and environmental effects.

Computational demand represents another practical consideration. For example, processing the rice and Scots pine datasets required less than five hours on the HPC infrastructure used in this study, although actual runtime depended on resource availability and cluster workload. WISER incorporates parallelized algorithms, including an ABC-based procedure for variance-component estimation, to facilitate analyses of complex experimental designs. Such approaches may provide practical advantages over some iterative variance-component estimation algorithms, which can suffer from slow or difficult convergence in complex settings [77–79].

Nevertheless, scalability is likely to become increasingly important for populations substantially larger than those investigated here. In particular, datasets exceeding approximately 100,000 observations may pose computational challenges under the current implementation. Future work should therefore evaluate WISER’s computational scalability and investigate strategies for extending the framework to substantially larger datasets.

### 4.7. Historical context, implications for breeding programs, versatility of the whitening approach, and future directions

Mixed-model approaches, particularly BLUP, have long provided a practical framework for BV estimation and remain widely used because of their statistical foundations, flexibility, and implementation in accessible software, including R packages such as `lme4` [62]. However, when genotype representation is strongly unbalanced or genetic structure is confounded with environmental effects, conventional modeling assumptions may not fully account for dependencies between the genetic and experimental components of the design.

WISER addresses this problem from a different perspective by using the genetic covariance structure to whiten the fixed-effect design before estimating BVs. The results presented here suggest that this approach can be particularly useful in structured and unbalanced experimental populations, providing more stable BVs and improving subsequent genomic prediction.

These properties are directly relevant to breeding programs, where BV estimates often constitute intermediate phenotypes used for subsequent genomic analyses and selection decisions. The smaller interquartile ranges of GBLUP PA observed with WISER suggest greater stability in predictive performance across cross-validation replicates. Such stability may be particularly valuable when genomic predictions are used to support selection decisions across heterogeneous experimental conditions.

Several extensions could further broaden the applicability of WISER. Incorporating additional environmental descriptors could improve the characterization of heterogeneous experimental conditions, while extending the framework to explicitly model genotype-by-environment interactions could improve BV estimation when genetic effects vary substantially among environments. Beyond the genomic prediction models evaluated in this study, future work could explore the integration of WISER with more complex predictive approaches, including deep learning-based models.

More broadly, the whitening principle underlying WISER provides a flexible way to account for genetic covariance during adjustment for environmental effects without directly estimating BVs from marker effects. This separation between BV estimation and downstream genomic prediction makes the framework applicable beyond the specific datasets and prediction models investigated here.

## 5. Conclusion

In conclusion, this study provides evidence that WISER can improve BV estimation, particularly in datasets characterized by genotype–environment confounding, unequal genotype representation, and complex genetic structure. Results from empirical datasets and simulations indicate that whitening the fixed-effect design can improve BV estimation and downstream genomic prediction. From an algebraic perspective, WISER can be interpreted as a continuous spectral reweighting of the fixed-effect design according to the eigenspectrum of the genetic covariance matrix. Rather than explicitly correcting for or removing a predefined subset of genetic eigendirections, WISER assigns relatively smaller weights to directions associated with larger eigenvalues, which represent dominant axes of genetic variation and may capture major components of population structure. WISER thereby preserves information across the full eigenspace while reducing the relative influence of its dominant directions. This spectral interpretation provides a plausible explanation for how WISER may reduce confounding between genetic structure and environmental effects. Importantly, the simulation results, marker-subsampling sensitivity analysis, and two-stage GBLUP comparison suggest that the benefits of WISER cannot be explained solely by repeated use of genomic information, but are also related to the whitening transformation itself. WISER therefore provides a complementary framework for BV estimation in structured and unbalanced experimental populations, with potential applications in large-scale crop and livestock breeding programs.

## Abbreviations

ABC: Approximate Bayesian computation
AIC: Akaike information criterion
ANOVA: Analysis of variance
BLUE: Best linear unbiased estimation
BLUP: Best linear unbiased prediction
CH: index Calinski-Harabasz index
CV: Cross-validation
D: Duration of budburst
EA: Estimation accuracy
EM: Expectation-maximization
EVG: covariance analysis via eigenvectors
FL: Days to flowering
GBLUP: Genomic best linear unbiased prediction
GPS: Global positioning system
GWAS: Genome-wide association studies H Height
I: Annual height increment
IQR: Interquartile range
LASSO: Least absolute shrinkage and selection operator
LS-means: Least squares means
NIRS: Near-infrared spectroscopy
OLS: Ordinary least squares
PA: Predictive ability
PC: Principal component
PCA: Principal component analysis
PCA-cor: Principal component correlation analysis
PH: Plant height
QTL: Quantitative trait locus
REFPOP: Reference population
REML: Restricted maximum likelihood
RF: Random forest
RKHS: Reproducing kernel Hilbert space
SNP: Single nucleotide polymorphism
SVR: Support vector regression
T4: Time taken to reach individual budburst stages: stage 4
T5: Time taken to reach individual budburst stages: stage 5
T6: Time taken to reach individual budburst stages: stage 6
UMAP: Uniform manifold approximation and projection
VIF: Variance inflation factor
WISER: Whitening and successive least squares estimation refinement
YLD: Grain yield
ZCA-cor: Zero-phase component correlation analysis Z
N: Zinc concentration

## Supplementary information

Supplementary File 1

Algebraic derivations which show the relationship between WISER and eigenvector-based population structure corrections.

Supplementary File 2

Presents a detailed overview of the ABC algorithm implemented within the WISER framework for variance components (i.e., 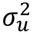 and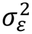 ) estimation, along with the regularization and shrinkage techniques used to ensure positive-definiteness for the genetic covariance matrix Σ_*u*_. Supplementary File 3

Presents median predictive ability (PA) values and interquartile ranges (IQR) in parentheses, across all species and traits, for several prediction models including random forest (RF), support vector regression (SVR), genomic best linear unbiased prediction (GBLUP), reproducing kernel Hilbert space (RKHS) regression, and least absolute shrinkage and selection operator (LASSO).

Supplementary File 4

Presents the completeness of the experimental designs used in this study. For the rice, maize, and Scots pine datasets, it reports the completeness of each block within each environment by indicating whether all genotypes observed at the corresponding site are represented in the block. For the apple REFPOP dataset, it reports the completeness of management groups within environments. These summaries were used to characterize the degree of imbalance in genotype representation across blocks, managements, environments, and sites, and to illustrate the incompleteness commonly encountered in practical breeding datasets that may contribute to confounding between genetic and environmental effects.

Supplementary File 5

Presents GBLUP median predictive ability (PA) values and interquartile ranges (IQR) obtained on the rice dataset when the whitening covariance matrix used by WISER was estimated from a random subset of only 100 SNPs, whereas genomic prediction was performed using the complete marker dataset (9,929 SNPs). This sensitivity analysis was conducted to assess the robustness of WISER to a substantial reduction in the marker information used to characterize population structure during the whitening step.

Supplementary File 6

Presents an evaluation of a two-stage GBLUP framework in which BVs estimated from a first-stage GBLUP model are subsequently used as response variables in a second GBLUP model incorporating genomic covariance information derived from marker data. Median predictive ability (PA) and genomic heritability (*h*^2^) values obtained for the rice and Scots pine datasets are reported and compared with those obtained using WISER, LS-means, BLUP, and BLUP-PCA BVs. The supplementary file also provides a theoretical analysis of the two-stage GBLUP framework, showing that its predictive performance remains fundamentally dependent on the quality of the BVs estimated during the first stage and explaining why repeated incorporation of genomic covariance information alone does not reproduce the gains observed with WISER.

Supplementary Figures 1–4

PCA and UMAP 2D projections of the apple REFPOP genomic data, shown with and without family cluster labels to visualize population structure.

Supplementary Figures 5–16

Heatmaps, PCA, and UMAP 2D plots illustrating population structure in the rice, maize, apple, and pine genomic datasets.

Supplementary Figures 17–20

Boxplots and violin plots of the GBLUP predictive ability (PA) values for traits of each species, for BVs estimated using WISER, LS-means, and BLUP approaches.

Supplementary Figures 21–28

Boxplots and violin plots of mean squared error (MSE) and estimation accuracy (EA) between simulated and estimated genetic values for WISER (with whitening of fixed-effect variables) and BLUE (without whitening), based on a generic trait simulated with moderate heritability (0.5) under each species’ experimental design.

Supplementary Figures 29–36

Boxplots and violin plots of mean squared error (MSE) and estimation accuracy (EA) between simulated and naïvely estimated genetic values for WISER (with whitening of fixed-effect variables) and BLUE (without whitening), based on a generic trait simulated with moderate heritability (0.5) under each species’ experimental design.

## Authors’ contributions

LJ derived the algebra for the analytical results, created the GitHub repositories, developed the associated R scripts, developed the WISER R package and conducted all dataset analyses. LJ also authored the manuscript. WG, ML, AP, MR, CED, FL, LL, MJA, TMH, and HM reviewed and approved the final version of the manuscript.

## Authors’ information

The authors declare no conflict of interests.

## Funding

Phenotypic data collection was partially supported by the Horizon 2020 Framework Program of the European Union under grant agreement No 817970 (project INVITE: “Innovations in plant variety testing in Europe to foster the introduction of new varieties better adapted to varying biotic and abiotic conditions and to more sustainable crop management practices”) and by the SusCrop Agrobiodiversity project Apple BIOME (“Microbiome and genomic analysis in apple germplasm towards broadening genetic resources to breed for resilient varieties”). LJ was supported by the ANR grant for the Apple BIOME project (ANR-22-SUSC-0001-05).

## Acknowledgments

The authors thank the field technicians and staff at INRAe IRHS and Experimental Unit (UE Horti), Angers, France, the Fruit Breeding Group at Agroscope in Waedenswil, Switzerland, and technical staff at all apple REFPOP sites for the maintenance of the orchards and phenotypic data collection.

## Availability of data and materials

All datasets and R scripts used in this study are publicly available through the following GitHub repositories. Archived versions of the repositories are also available on Zenodo through their corresponding DOIs:

- Rice genomic prediction: [GitHub repository](https://github.com/ljacquin/wiser_genomic_prediction_rice); [Zenodo archive](https://doi.org/10.5281/zenodo.23345152)
- Maize genomic prediction: [GitHub repository](https://github.com/ljacquin/wiser_genomic_prediction_rice); [Zenodo archive](https://doi.org/10.5281/zenodo.23345152)
- Apple genomic prediction (refpop): [GitHub repository](https://github.com/ljacquin/wiser_genomic_prediction_rice); [Zenodo archive](https://doi.org/10.5281/zenodo.23345152)
- Scots pine genomic prediction: [GitHub repository](https://github.com/ljacquin/wiser_genomic_prediction_rice); [Zenodo archive](https://doi.org/10.5281/zenodo.23345152)
- Statistical analyses of WISER results: [GitHub repository](https://github.com/ljacquin/wiser_genomic_prediction_rice); [Zenodo archive](https://doi.org/10.5281/zenodo.23345152)
- Rice marker-subsampling sensitivity analysis: [GitHub repository](https://github.com/ljacquin/wiser_genomic_prediction_rice); [Zenodo archive](https://doi.org/10.5281/zenodo.23345152)
- Two-stage GBLUP analyses: [GitHub repository](https://github.com/ljacquin/wiser_genomic_prediction_rice); [Zenodo archive](https://doi.org/10.5281/zenodo.23345152)

## Declarations

### Ethics approval and consent to participate

Not applicable.

### Consent for publication

Not applicable.

### Competing interests

The authors declare no competing interests.

## Notes

https://doi.org/10.5281/zenodo.23344278

https://doi.org/10.5281/zenodo.23344326

https://doi.org/10.5281/zenodo.23344483

https://doi.org/10.5281/zenodo.23345121

https://doi.org/10.5281/zenodo.23345139

https://doi.org/10.5281/zenodo.23345152

https://doi.org/10.5281/zenodo.23345214

https://github.com/ljacquin/compute_stats_wiser_results

https://github.com/ljacquin/wiser

### Competing Interest Statement

The authors have declared no competing interest.

### Summary of Updates

This version of the manuscript has been revised to include new simulation studies evaluating WISER using both predictive ability (PA) and true predictive ability (TPA) criteria. These additional analyses provide a more comprehensive assessment of method performance across different validation scenarios and levels of confounding between genetic and environmental effects. The simulation section, associated results, figures, and supplementary materials have been updated accordingly. The new results further characterize the conditions under which WISER improves prediction accuracy relative to conventional approaches and provide additional insight into the relationship between estimated and true predictive performance. Minor revisions were also made throughout the manuscript to improve clarity, consistency, and interpretation of the simulation results.

