## Supplementary File 1 for "WISER: spectral whitening of fixed effects for improved breeding value estimation in structured, unbalanced, and genotype–environment-confounded populations"

**Algebraic relationship between WISER and eigenvector-based population structure corrections:**

Similar to $R_{u}$ (see Section 2.2), the spectral decomposition of $\Sigma_{u}$ can be expressed as $\Sigma_{u}= U\Lambda U'$, where $\Lambda\in\mathbb{R}^{r\times r}$ is the diagonal matrix of $r$ positive eigenvalues, and 𝑈$\in\mathbb{R}^{n\times r}$ the matrix whose columns are the orthonormal eigenvectors associated with the positive eigenvalues of $\Sigma_{u}$. Note that $rank\left( \Sigma_{u} \right)=rank\left( ZKZ' \right)= rank\left( ZPDP'Z' \right)=rank\left( (ZP)D(ZP)' \right)$where $D$ is the eigenvalue matrix of $K$, hence $rank\left( \Sigma_{u} \right)=r\leq rank\left( K \right)\leq q<n$ , $U^{'}U=I_{r}$ and $UU^{'}\neq I_{n}$. If $K$ is full rank ($rank\left( K \right)=q$) and the incidence matrix $Z$ does not reduce the rank of $ZP$, then $rank\left( \Sigma_{u} \right)=r=q.$

According to [6], the EVG model which accounts for population structure correction is defined as follows:

$$Y=X\beta+ \sum_{s=1}^{m} U_{s}\alpha_{s}+ Zu+\varepsilon(EVG)$$

where $\left( U_{s} \right)_{1\leq s\leq m}$represents the first $m\leq r$ chosen eigenvectors, with the highest eigenvalues, and $\left( \alpha_{s} \right)_{1\leq s\leq m}$ their corresponding fixed effects. To align the notations of this model with those of the WISER approach, particularly with respect to the spectral decomposition of $\Sigma_{u}$, we assume all $r$ eigenvectors associated with the positive eigenvalues of $\Sigma_{u}$ are included in the model. In this case, the EVG model can be rewritten as:

$$Y=X\beta+ U\alpha+\xi(EVG)$$

where $U$ is the eigenvector matrix of $\Sigma_{u}$, $\alpha=\left[ \alpha_{1}, \ldots,\alpha_{r} \right]^{'}$ and $\xi=Zu+\varepsilon.$ As implicitly stated in Section 2.2, WISER also assumes that $\xi=Zu+\varepsilon$, but within its framework, the vector $u$ is replaced by $v$, for which no distributional assumption is made. This allows $v$to be estimated without explicitly expressing it as a linear or nonlinear combination of omic effects.

Let$E_{U}=span\left( U \right)$ represent the vector subspace generated by the column vectors of $U$. Let $P_{E_{U}}=U\left( U^{'}U \right)^{-1}U^{'}=UU'$ and $P_{E_{U}^{\perp}}=I_{n}-P_{E_{U}}=I_{n}-UU^{'}$ be the orthogonal projectors onto $E_{U}$ and $E_{U}^{\perp}$ respectively, where $E_{U}^{\perp}$is the orthogonal complement of $E_{U}$ (i.e., $\mathbb{R}^{n}=E_{U}\oplus E_{U}^{\perp}$). Using the following linear transformation based on $E_{U}^{\perp}$, we have:

$$Y=X\beta+ U\alpha+\xi\left( EVG \right)$$

$$\Leftrightarrow P_{E_{U}^{\perp}}Y=P_{E_{U}^{\perp}}X\beta+\underset{=0}{\underbrace{P_{E_{U}^{\perp}}U}\alpha}+P_{E_{U}^{\perp}}\xi$$

$$\Leftrightarrow Y^{*}= X^{*}\beta+ \xi^{*}$$

where $Y^{*}= P_{E_{U}^{\perp}}Y$, $X^{*}= P_{E_{U}^{\perp}}X$ and $\xi^{*}= P_{E_{U}^{\perp}}\xi$. Note that $\left( U_{s} \right)_{1\leq s\leq r}\in$ $E_{U}$, therefore we have $P_{E_{U}^{\perp}}U=0$. The last property can also be verified directly as follows:

$$P_{E_{U}^{\perp}}U={(I}_{n}-UU^{'})U=U-UU^{'}U=U-U=0$$

In the absence of assumptions regarding $\xi^{*}$, as is the case within the WISER framework, and provided that $P_{E_{U}^{\perp}}X$has full column rank, the OLS estimate for $\beta$ in the $EVG$ model is given by:

$$\hat{\beta}_{EVG}= \left( X^{*}'X^{*} \right)^{-1}X^{*}'Y^{*}=\left( {(P}_{E_{U}^{\perp}}{X)'P}_{E_{U}^{\perp}}X \right)^{-1}{(P}_{E_{U}^{\perp}}X)'P_{E_{U}^{\perp}}Y$$

$$\Leftrightarrow\hat{\beta}_{EVG}=\left( X'P_{E_{U}^{\perp}}{'P}_{E_{U}^{\perp}}X \right)^{-1}X'P_{E_{U}^{\perp}}'P_{E_{U}^{\perp}}Y$$

$$\Leftrightarrow\hat{\beta}_{EVG}= \left( X'P_{E_{U}^{\perp}}X \right)^{-1}X'P_{E_{U}^{\perp}}Y$$

Note that $P_{E_{U}^{\perp}}$ is symmetric and idempotent, therefore we have $P_{E_{U}^{\perp}}{'P}_{E_{U}^{\perp}}$= $P_{E_{U}^{\perp}}P_{E_{U}^{\perp}}$= $P_{E_{U}^{\perp}}.$ From the expression for $\hat{\beta}_{EVG}$, one can notice that the vectors of the design matrix $X$ for fixed effects are orthogonally projected onto $E_{U}^{\perp}$ (i.e., $P_{E_{U}^{\perp}}X$), which represents the orthogonal complement of $E_{U}$. This projection serves to reduce the alignment between fixed effects and the principal directions of genetic variation, thereby potentially attenuating genetic-environment confounding. Indeed, the fixed-effect design matrix can be decomposed as follows:

$$X=UA+R$$

with $A=U'X$ and $R=\left( I-UU^{'} \right)X$, where $UA=P_{E_{U}}X$ represents the component of the fixed-effect design matrix lying in the eigenspace of the genetic covariance matrix, and $R$ the component orthogonal to this eigenspace. Hence, we have $P_{E_{U}^{\perp}}X=P_{E_{U}^{\perp}}\left( UA+R \right)=R,$ which shows that the EVG correction completely removes the component of the fixed-effect design matrix lying in the eigenspace of the genetic covariance matrix while preserving its orthogonal component. However, in practice, only a subset of eigenvectors is typically included in the EVG model. Consequently, only a part of the genetic covariance structure is accounted for, and some alignment between fixed effects and genetic variation may remain.

Furthermore, the EVG correction can be expressed within the same transformed-OLS framework as WISER. Specifically, the projector $P_{E_{U}^{\perp}}$may be viewed as a pseudo-whitening operator in an algebraic sense, since substituting it for the whitening matrix in the WISER OLS estimator recovers the EVG estimator. Recall that the WISER estimator of $\beta$can be written as:

$$\hat{\beta}_{WISER}=\left( \tilde{X}'\tilde{X} \right)^{-1}\tilde{X}'Y=\left( X'W'WX \right)^{-1}X'W'Y$$

Let

$$W_{Pseudo}=P_{E_{U}^{\perp}}$$

Since $P_{E_{U}^{\perp}}$ is symmetric and idempotent, we have:

$$\begin{aligned} \hat{\beta}_{Pseudo WISER}&=\left( X^{'}{W'}_{Pseudo}W_{Pseudo}X \right)^{-1}X^{'}{W'}_{Pseudo}Y \\ &=\left( X^{'}P_{E_{U}^{\perp}}^{'}P_{E_{U}^{\perp}}X \right)^{-1}X^{'}P_{E_{U}^{\perp}}^{'}Y \\ &=\left( X^{'}P_{E_{U}^{\perp}}X \right)^{-1}X^{'}P_{E_{U}^{\perp}}Y \\ &=\hat{\beta}_{EVG} \end{aligned}$$

provided that $P_{E_{U}^{\perp}}X$has full column rank. Thus, at the level of fixed-effect estimation, the EVG correction can be represented as a particular transformed-OLS formulation in which $P_{E_{U}^{\perp}}$replaces the whitening operator used by WISER.

However, $W_{Pseudo}$ is not a whitening operator in the strict covariance sense. For $T=Zu$, with

$$Var\left( T \right)=\Sigma_{u}=U\Lambda U^{'}$$

we obtain:

$$\begin{aligned} Var\left( W_{Pseudo}T \right)&=W_{Pseudo}\Sigma_{u}{W'}_{Pseudo} \\ &=P_{E_{U}^{\perp}}\left( U\Lambda U^{'} \right)P_{E_{U}^{\perp}} \\ &=0 \end{aligned}$$

because $P_{E_{U}^{\perp}}U=0$. Therefore, $P_{E_{U}^{\perp}}$does not whiten the genetic covariance in the strict sense. Instead, it completely removes the variation associated with the genetic eigendirections contained in $E_{U}$. WISER differs fundamentally in that it does not remove these directions but reweights their contribution according to their associated eigenvalues. Thus, EVG can be viewed as a hard spectral correction, whereas WISER performs a continuous spectral reweighting of the genetic eigendirections. To illustrate specifically how whitening reweights the component of $X$ lying in $E_{U}$, consider the spectral operator:

$$W_{E_{U}}=U\Lambda^{-1/2}U^{'}$$

restricted here to the eigenspace of $\Sigma_{u}$. Its action on $UA$ is:

$$W_{E_{U}}(UA)=U\Lambda^{-\frac{1}{2}}A=\sum_{i=1}^{r} \frac{1}{\surd\lambda_{i}}U_{i}A_{i}$$

where $U_{i}$ is the $i$-th eigenvector of $\Sigma_{u}$ (a column vector of dimension $n\times1$), and $A_{i}$ is the $i$-th row of $A$(a row vector of dimension $1\times l$) containing the coefficients of the fixed-effect design matrix along the $i$-th genetic eigendirection. Hence, this spectral reweighting transformation rescales the contribution of each eigendirection according to the factor $\lambda_{i}^{-1/2}$. Directions associated with large eigenvalues, which correspond to dominant axes of genetic variation and may capture major components of population structure, are therefore assigned smaller weights, whereas directions associated with smaller eigenvalues are relatively less affected. Consequently, the component

$$UA=\sum_{i=1}^{r} U_{i}A_{i}$$

is transformed into

$$U\Lambda^{-\frac{1}{2}}A=\sum_{i=1}^{r} \frac{1}{\surd\lambda_{i}}U_{i}A_{i}$$

so that each eigendirection is reweighted according to its associated eigenvalue. Unlike EVG, which completely removes the component $UA$, the spectral action of WISER retains the contribution of these eigendirections while differentially reweighting them according to their associated eigenvalues. Therefore, WISER may be interpreted as a softer correction strategy that attenuates dominant directions of genetic variation, including those associated with population structure, while retaining their contribution to the transformed fixed-effect design. Note that the operator $U\Lambda^{-\frac{1}{2}} U^{'}$ is introduced here solely to illustrate the spectral reweighting induced by whitening.

In the actual WISER framework, the whitening property is ensured through the regularization of $\Sigma_{u}$, yielding a full-rank covariance matrix that prevents the over-amplification of components associated with near-zero eigenvalues. Indeed, in the WISER framework the regularized form of $\Sigma_{u}$is given by:

$$\tilde{\Sigma}_{u}=\Sigma_{u}+\alpha\sigma\left( \Sigma_{u} \right)I_{n}$$

where $\alpha>0$ is a regularization parameter and $\sigma\left( \Sigma_{u} \right)>0$ is a scalar measure of the magnitude of $\Sigma_{u}$. The eigenvalues associated with the non-null eigendirections become

$$\lambda_{i}+\alpha\sigma\left( \Sigma_{u} \right),$$

whereas directions belonging to the null space of $\Sigma_{u}$ receive the strictly positive eigenvalue

$$\alpha\sigma\left( \Sigma_{u} \right)$$

Consequently, all eigendirections are assigned positive eigenvalues, preventing the excessive amplification that would otherwise arise for directions associated with near-zero eigenvalues under standard whitening. In the datasets considered here, $\alpha\sigma\left( \Sigma_{u} \right)$ was consistently greater than 1 for $\alpha\in\left[ 0.1 , 1 \right]$, further limiting the influence of weakly supported eigendirections.

The relationship between the EVG and principal component (PC) models can be demonstrated as follows. According to [6], the PC model, which corrects for population structure, is defined as follows:

$$Y=X\beta+ Q\gamma+\xi\left( PC \right)$$

where $Q$ contains the coordinates of individuals along principal components derived from the genetic covariance matrix. Because the principal components are derived from eigendirections of $\Sigma_{u}$, the columns of $Q$belong to $E_{U}=\mathrm{span}\left( U \right)$. Therefore,

$$\mathrm{col}(Q)\subseteq E_{U}$$

which implies

$$P_{E_{U}^{\perp}}Q=0$$

Consequently, under the assumption that all retained eigendirections are included, applying the orthogonal projection $P_{E_{U}^{\perp}}$ to the PC model yields the same OLS estimate of the fixed effects as in the EVG formulation. Therefore, EVG and PC corrections may both be interpreted as approaches that account for population structure through selected eigen-information derived from the genetic covariance matrix.
