## Supplementary File 2 for "WISER: spectral whitening of fixed effects for improved breeding value estimation in structured, unbalanced, and genotype–environment-confounded populations"

**WISER approximate Bayesian computation (ABC) algorithm:**

The ABC algorithm implemented in WISER for estimating $\sigma_{u}^{2}$ and $\sigma_{\varepsilon}^{2}$ is structured as follows:

1. Define the prior distributions for $\sigma_{u}^{2}$ and $\sigma_{\varepsilon}^{2}$:

$$\sigma_{u}^{2}\sim U({\sigma_{u, min}^{2},\sigma}_{u, max}^{2})$$

$$\sigma_{\varepsilon}^{2}\sim U({\sigma_{\varepsilon, min}^{2},\sigma}_{\varepsilon, max}^{2})$$

2. Simulate the data :

For each simulation $\left( i \right)$ (where $i=1,\ldots,N)$,

- Sample $\sigma_{u}^{2, (i)}$ and $\sigma_{\varepsilon}^{2,(i)}$ from their prior distributions
- Sample the random genetic effects: $u^{(i)} \sim N_{q}(0, \sigma_{u}^{2, (i)}K)$
- Sample the random residuals: $\varepsilon^{(i)} \sim N_{n}(0, \sigma_{\varepsilon}^{2,(i)}I_{n})$
- Simulate the vector of phenotypes: $Y^{(i)}=X\hat{\beta}+Zu^{(i)}+\varepsilon^{(i)}$

3. Compute the distance $d^{(i)}$ between the observed and simulated phenotypes:

$d^{(i)}=$ $||Y-Y^{(i)}||_{2}^{2}$

4. Define an acceptance/rejection threshold $\delta$ for distances based on a $\alpha$-quantile:

$\delta= \alpha$-quantile ($d^{(1)},$.., $d^{(i)},..,d^{(N)})$

$\delta$ is the value estimated from the ordered distribution of $d^{\left( i \right)}$, such that the empirical distribution function captures $\alpha$% of the ordered distribution. In other words, only the distances that are less than or equal to $\delta$ are retained, representing $\alpha$% of the total.

5. Accept parameter sets :

Only parameter sets ${\{\sigma}_{u}^{2}, \sigma_{\varepsilon}^{2}{\}}^{(i)}$ for which $d^{(i)}\leq$ $\delta$ are accepted.

6. Estimate the parameters a posteriori:

After obtaining the distribution of distances, the a posteriori means of the parameters are computed as follows:

$$\hat{\sigma}_{u}^{2}=\frac{1}{M}\sum_{j=1}^{M} \sigma_{u}^{2,(j)}$$

$$\hat{\sigma}_{\varepsilon}^{2}=\frac{1}{M}\sum_{j=1}^{M} \sigma_{\varepsilon}^{2,(j)}$$

where $j=1,..,M$ corresponds to the indices of the accepted parameter sets.

For this algorithm, we chose the following parameters:

- $\sigma_{u, min}^{2}= \sigma_{\varepsilon, min}^{2}={10}^{-2}$, and $\sigma_{u, max}^{2}= \sigma_{\varepsilon, max}^{2}=\sigma_{y}^{2}$
- $N=100$
- $\alpha=0.05$

It is noted that this algorithm also requires the vector $\hat{\beta}$ in order to estimate $\sigma_{u}^{2}$ and $\sigma_{\varepsilon}^{2}$. Before the simulation step, this vector is calculated by setting $\sigma_{u}^{2}=\sigma_{\varepsilon}^{2}=1$. It is then recalculated after each estimation of the components $\sigma_{u}^{2}$ and $\sigma_{\varepsilon}^{2}$ by the proposed ABC algorithm, and this process is repeated at least once, or as many times as necessary, as defined by the user.

**WISER shrinkage and regularization procedures:**

Let $\tilde{\Sigma}_{u}^{1}$ denote a regularized and positive-definite form of $\Sigma_{u}$. A general and established approach for computing $\tilde{\Sigma}_{u}^{1}$ is given by [51, 53]:

$$\tilde{\Sigma}_{u}^{1} =\left( 1-\alpha\right)\Sigma_{u}+\alpha\sigma(\Sigma_{u})I_{n}$$

where $\alpha\in]0,1]$ is the shrinkage intensity parameter and $\sigma\left( \Sigma_{u} \right)>0$ is a scalar measure of the magnitude of $\Sigma_{u}$ which scales with its dimensions. The computed matrix $\tilde{\Sigma}_{u}^{1}$ is generally referred to as a shrinkage estimator because it progressively shrinks $\left( 1-\alpha\right)\Sigma_{u}$ as $\alpha$ increases, converging toward $\alpha\sigma(\Sigma_{u})I_{n}$, a matrix proportional to the identity matrix. Possible choices for $\sigma\left( \Sigma_{u} \right)$ include the trace, $\sigma\left( \Sigma_{u} \right)=tr\left( \Sigma_{u} \right)$, or any matrix norm, such as the Frobenius norm, $\sigma\left( \Sigma_{u} \right)= \parallel\Sigma_{u}\parallel_{F} = \sqrt{tr(\Sigma_{u}'\Sigma_{u})}$, which are both implemented in WISER.

However, as $\alpha$ increases, both the diagonal and off-diagonal elements of the covariance structure associated to $\Sigma_{u}$ are modified. This alteration may be undesirable, especially when the off-diagonal elements, which capture essential covariance information, are critical. To avoid such modifications, a simpler regularized and positive-definite form of $\Sigma_{u}$ can be used:

$$\tilde{\Sigma}_{u}^{2}=\Sigma_{u}+\alpha\sigma(\Sigma_{u})I_{n}$$

In WISER, both $\tilde{\Sigma}_{u}^{1}$ and $\tilde{\Sigma}_{u}^{2}$ are available, with $\tilde{\Sigma}_{u}^{2}$and $\sigma\left( \Sigma_{u} \right)= \parallel\Sigma_{u}\parallel_{F}$ used by default. The shrinkage intensity parameter $\alpha$ can be determined via K-fold cross-validation to minimize mean square error (MSE) based on omics data. The predictive models implemented for this purpose in WISER include random forest (RF), support vector regression (SVR), the best linear unbiased predictor (BLUP), reproducing kernel Hilbert space (RKHS) regression, and the least absolute shrinkage selection operator (LASSO).

For simulation purposes, to maintain computational tractability while preserving a comparable level of regularization across sample sizes, the diagonal regularization parameter was defined as:

$$\alpha={\alpha_{ref}\left( \frac{n_{ref}}{n} \right)}^{\frac{1}{2}}$$

where$\alpha_{ref}=1$, $n_{ref}=1000$, and $n$denotes the number of phenotypic observations (i.e., the dimension of $\Sigma_{u}$). Regularization was applied by adding $\alpha\parallel\Sigma_{u}\parallel_{F}I$to $\Sigma_{u}$, where $\parallel\Sigma_{u}\parallel_{F}$denotes the Frobenius norm. Because $\parallel\Sigma_{u}\parallel_{F}$increases with matrix dimension, the magnitude of the added diagonal term depends on sample size. More generally, the Frobenius norm scales approximately as a power of the matrix dimension:

$$\parallel\Sigma_{u}\parallel_{F}\approx kn^{\gamma}$$

where $k$is a constant determined by the variance scale and $\gamma$typically ranges from 1/2 for nearly diagonal matrices to $1$for dense matrices. For the covariance structures considered in this study, empirical examination indicated that $\parallel\Sigma_{u}\parallel_{F}$scaled approximately as $kn^{1/2}$. Consequently, enforcing invariance of the added diagonal penalty relative to a reference sample size $n_{ref}$ yields:

$$\alpha\left( n \right)\parallel\Sigma_{u}\left( n \right)\parallel_{F}= \alpha\left( n_{ref} \right)\parallel\Sigma_{u}\left( n_{ref} \right)\parallel_{F}$$

$$\Leftrightarrow\alpha\left( n \right)kn^{1/2}= \alpha\left( n_{ref} \right)kn_{ref}^{1/2}$$

$$\Leftrightarrow\alpha={\alpha_{ref}\left( \frac{n_{ref}}{n} \right)}^{\frac{1}{2}}$$

This scaling was intended to maintain a broadly comparable level of regularization across simulation settings. The reference value was calibrated at $n_{ref}=1000$, for which estimation accuracy and predictive ability relative to the true genetic values were stable over the range $0.1\leq\alpha\leq1$.
