## Supplementary File 5 for "WISER: spectral whitening of fixed effects for improved breeding value estimation in structured, unbalanced, and genotype–environment-confounded populations"

**Evaluating WISER under reduced marker information: whitening based on a random subset of 100 SNPs**

Table S5 – GBLUP median PA values for BV estimation methods (WISER, LS-means, and BLUP approaches) across all traits in the rice dataset, with interquartile ranges (IQR) in parentheses. Values are based on 100 cross-validation (CV) scenarios.

| Dataset | Trait | GBLUP median PA values | | | |
| --- | --- | --- | --- | --- | --- |
|  |  | WISER | LS-means | BLUP | BLUP-PCA |
| Rice dataset | days to flowering (FL) | 0.61 (0.10) | 0.41 (0.12) | 0.41 (0.11) | 0.21 (0.11) |
| 334 genotypes | plant height (PH) | 0.61 (0.09) | 0.51 (0.09) | 0.51 (0.09) | 0.42 (0.09) |
| 9,929 SNPs | grain yield (YLD) | 0.47 (0.10) | 0.49 (0.11) | 0.49 (0.11) | 0.42 (0.10) |
|  | grain zinc concentration (ZN) | 0.52 (0.11) | 0.36 (0.16) | 0.36 (0.16) | 0.26 (0.19) |
