## Supplementary File 6 for "WISER: spectral whitening of fixed effects for improved breeding value estimation in structured, unbalanced, and genotype–environment-confounded populations"

**Evaluation of a two-stage GBLUP framework and comparison with WISER**

To further investigate whether the improvements obtained with WISER could arise from a repeated use of genomic covariance information across successive analytical steps, an additional two-stage GBLUP analysis was performed on the rice and Scots pine datasets. These datasets were selected because their size made the additional computations tractable within a reasonable time frame.

The two-stage GBLUP framework consisted of the following successive models:

$$M_{1}: Y=X\beta+Z_{u_{1}}u_{1}+\varepsilon_{1},$$

with

$$u_{1}\sim N_{q}\left( 0,\sigma_{g}^{2}K \right),\varepsilon_{1}\sim N_{n}\left( 0,\sigma_{\varepsilon_{1}}^{2}I_{n} \right),$$

followed by

$$M_{2}: \hat{u}_{1}=\mu+Z_{u_{2}}u_{2}+\varepsilon_{2},$$

with

$$u_{2}\sim N_{q}\left( 0,\sigma_{g}^{2}K \right), \varepsilon_{2}\sim N_{q}\left( 0,\sigma_{\varepsilon_{2}}^{2}I_{q} \right).$$

In this framework, breeding values estimated in the first GBLUP step ($M_{1}$), denoted $\hat{u}_{1}$ and hereafter referred to as GBLUP-BVs, were subsequently used as response variables in a second GBLUP model ($M_{2}$) that again incorporated genomic covariance information derived from marker data. This analysis was motivated by the hypothesis that the use of genomic covariance information during BV estimation, followed by its reuse during downstream GBLUP prediction, could by itself account for part of the gains observed with WISER. Under this hypothesis, GBLUP-BVs, for which genomic covariance information is incorporated at both stages, would be expected to show a substantial advantage over conventional BLUP BVs in downstream PA and $h^{2}$.

However, the results presented in Tables S6.1 and S6.2 show that the PA and $h^{2}$estimates obtained with GBLUP-BVs were very similar to those obtained with BLUP breeding values. In most cases, differences between BLUP and GBLUP-BVs were negligible. For example, in the rice dataset, the median PAs obtained with GBLUP-BVs were identical to those obtained using BLUP for all four traits. A similar pattern was observed in the Scots pine dataset, with only minor differences in PA and $h^{2}$ estimates between BLUP and GBLUP-BVs. By contrast, WISER consistently produced substantially higher PA and $h^{2}$ estimates.

These results indicate that incorporating genomic covariance information during BV estimation and reusing it during downstream GBLUP prediction is not, by itself, sufficient to reproduce the gains observed with WISER. Consequently, these gains are unlikely to result solely from repeated use of genomic covariance information across analytical stages.

Table S6.1 – GBLUP median PA values for BV estimation methods (WISER, LS-means, BLUP approaches, and GBLUP-BVs) across all traits of the rice and pine datasets, with interquartile ranges (IQR) in parentheses. Values are based on 100 cross-validation (CV) scenarios.

| Dataset | Trait | GBLUP median PA values | | | |  |
| --- | --- | --- | --- | --- | --- | --- |
|  |  | WISER | LS-means | BLUP | BLUP-PCA | GBLUP-BVs |
| Rice dataset | days to flowering (FL) | 0.74 (0.05) | 0.41 (0.12) | 0.41 (0.11) | 0.21 (0.11) | 0.41 (0.12) |
| 334 genotypes | plant height (PH) | 0.73 (0.08) | 0.51 (0.09) | 0.51 (0.09) | 0.42 (0.09) | 0.51 (0.09) |
| 9,929 SNPs | grain yield (YLD) | 0.62 (0.08) | 0.49 (0.11) | 0.49 (0.11) | 0.42 (0.10) | 0.49 (0.11) |
|  | grain zinc concentration (ZN) | 0.69 (0.05) | 0.36 (0.16) | 0.36 (0.16) | 0.26 (0.19) | 0.36 (0.16) |
| Pine dataset | D | 0.66 (0.10) | 0.08 (0.19) | 0.14 (0.17) | 0.11 (0.19) | 0.16 (0.18) |
| 208 genotypes | H | 0.72 (0.08) | 0.18 (0.17) | 0.31 (0.16) | 0.23 (0.18) | 0.30 (0.15) |
| 20,796 SNPs | I | 0.35 (0.25) | 0.10 (0.21) | 0.31 (0.20) | 0.20 (0.20) | 0.32 (0.22) |
|  | T4 | 0.39 (0.22) | 0.24 (0.18) | 0.32 (0.20) | 0.30 (0.23) | 0.34 (0.21) |
|  | T5 | 0.26 (0.25) | 0.14 (0.22) | 0.26 (0.24) | 0.12 (0.24) | 0.26 (0.26) |
|  | T6 | 0.20 (0.27) | 0.14 (0.24) | 0.20 (0.26) | 0.11 (0.28) | 0.20 (0.26) |

Table S6.2 – GBLUP median $h^{2}$ values for BV estimation methods (WISER, LS-means, BLUP approaches, and GBLUP-BVs) across all species and traits, with interquartile ranges (IQR) in parentheses. Values are based on 100 cross-validation (CV) scenarios.

| Dataset | Trait | GBLUP median $h^{2}$ values | | | |  |
| --- | --- | --- | --- | --- | --- | --- |
|  |  | WISER | LS-means | BLUP | BLUP-PCA | GBLUP-BVs |
| Rice dataset | days to flowering (FL) | 0.87 (0.02) | 0.53 (0.06) | 0.53 (0.06) | 0.00 (0.00) | 0.53 (0.06) |
| 334 genotypes | plant height (PH) | 0.86 (0.03) | 0.76 (0.03) | 0.76 (0.03) | 0.70 (0.05) | 0.76 (0.04) |
| 9,929 SNPs | grain yield (YLD) | 0.82 (0.04) | 0.73 (0.05) | 0.73 (0.05) | 0.68 (0.06) | 0.73 (0.05) |
|  | grain zinc concentration (ZN) | 0.84 (0.03) | 0.55 (0.10) | 0.56 (0.10) | 0.42 (0.17) | 0.55 (0.10) |
| Pine dataset | D | 0.95 (0.03) | 0.00 (0.17) | 0.03 (0.22) | 0.00 (0.07) | 0.20 (0.33) |
| 208 genotypes | H | 0.97 (0.01) | 0.30 (0.39) | 0.56 (0.16) | 0.44 (0.52) | 0.56 (0.15) |
| 20,796 SNPs | I | 0.73 (0.16) | 0.00 (0.01) | 0.62 (0.22) | 0.00 (0.00) | 0.65 (0.22) |
|  | T4 | 0.74 (0.14) | 0.54 (0.22) | 0.62 (0.22) | 0.59 (0.24) | 0.64 (0.22) |
|  | T5 | 0.52 (0.23) | 0.30 (0.44) | 0.48 (0.28) | 0.00 (0.23) | 0.47 (0.30) |
|  | T6 | 0.28 (0.34) | 0.08 (0.31) | 0.30 (0.35) | 0.00 (0.11) | 0.32 (0.32) |

The behavior observed in Tables S6.1 and S6.2 can be interpreted from the relationship between the response and the genetic effect in the second-stage GBLUP.

Since $Z_{u_{2}}=I_{q}$, model $M_{2}$ can be rewritten as

$$\hat{u}_{1}=\mu+u_{2}+\varepsilon_{2}.$$

Under the model assumptions,

$$\mathrm{Cov}(u_{2},\varepsilon_{2})=0.$$

Consequently,

$$\mathrm{Cov}(\hat{u}_{1},u_{2})=\mathrm{Cov}(u_{2}+\varepsilon_{2},u_{2})=\mathrm{Var}(u_{2})=\sigma_{g}^{2}K.$$

Similarly,

$$\mathrm{Var}\left( \hat{u}_{1} \right)=\mathrm{Var}\left( u_{2} \right)+\mathrm{Var}(\varepsilon_{2})=\sigma_{g}^{2}K+\sigma_{\varepsilon_{2}}^{2}I_{q}.$$

For genotype $i$, the correlation between $\hat{u}_{1i}$ and $u_{2i}$is therefore

$$\mathrm{Corr}(\hat{u}_{1i},u_{2i})= \frac{\mathrm{Cov}\left( \hat{u}_{1i},u_{2i} \right)}{\sqrt{Var(\hat{u}_{1i})}\sqrt{Var(u_{2i})}}=\frac{\sigma_{g}^{2}K_{ii}}{\sqrt{\sigma_{g}^{2}K_{ii}+\sigma_{\varepsilon_{2}}^{2}}\sqrt{\sigma_{g}^{2}K_{ii}}}=\sqrt{\frac{\sigma_{g}^{2}K_{ii}}{\sigma_{g}^{2}K_{ii}+\sigma_{\varepsilon_{2}}^{2}}}.$$

If $K$ is normalized such that $K_{ii}=1$, this expression simplifies to

$$\mathrm{Corr}(\hat{u}_{1i},u_{2i})=\sqrt{\frac{\sigma_{g}^{2}}{\sigma_{g}^{2}+\sigma_{\varepsilon_{2}}^{2}}}.$$

This expression illustrates that the correlation between the response and the genetic effect in the second-stage model depends on the relative contribution of the genomic and residual variance components. As $\sigma_{\varepsilon_{2}}^{2}$ increases relative to $\sigma_{g}^{2}$, this correlation decreases. A large $\sigma_{\varepsilon_{2}}^{2}$ indicates that a substantial proportion of the variation in the first-stage BVs is not captured by the genomic covariance structure specified through $K$. This residual variation may reflect estimation noise, model misspecification, or other sources of variation not adequately represented by $K$. It may also partly arise from an imperfect separation of genetic and environmental effects during first-stage BV estimation, although $\sigma_{\varepsilon_{2}}^{2}$ should not itself be interpreted as a direct measure of genotype–environment confounding. Consequently, reapplying genomic covariance information to first-stage BVs does not necessarily recover genetic signal that is weakly represented in these initial estimates. This interpretation is consistent with the limited differences between BLUP and GBLUP-BVs observed in Tables S6.1 and S6.2 and contrasts with WISER, which uses genomic covariance information to whiten the fixed-effect design before BVs are estimated.
